# Structural basis for autoinhibition and self-activation in the *Pseudomonas aeruginosa* virulence factor protease IV (PIV)

**DOI:** 10.64898/2026.09.07.749839

**Authors:** Christopher J. Harding, Carlin Hamill, Qingtian Gong, Clarissa Melo Czekster

## Abstract

Secreted proteases enable *Pseudomonas aeruginosa* to damage host tissues and evade host immune defences, but the molecular basis by which protease IV (PIV) is maintained as an inactive precursor remains unclear. Here we determined a 2.17 Å crystal structure of catalytically inactive PIV_S409A_. The precursor comprises an N-terminal Cap region, a CUB domain and a C-terminal trypsin-like protease domain. The Cap region forms an extended clamp across the precursor, and residues at the Cap–CUB junction lie above the catalytic cleft. Structural comparison with a modelled peptide substrate indicates that the Cap does not engage the substrate-recognition pockets as a pseudosubstrate; instead, it sterically prevents access to the S1 pocket, oxyanion hole and catalytic triad. Wild-type PIV undergoes progressive processing to a 27.6-kDa mature species, whereas the S409A variant remains resistant to activation by exogenous mature enzyme, supporting an intramolecular initiating step. Mature PIV is weakly inhibited by an ornithine-containing peptide mimicking the Cap domain, is not inhibited by phenylmethylsulfonyl fluoride and is slowly irreversibly inhibited by TLCK, demonstrating unique characteristics in comparison to other proteases from the same family. Finally, mature PIV increases the activity of aminopeptidase PaAP, likely through maturation of this enzyme, as part of a protease activation cascade. These findings define the architecture of the PIV precursor and provide a structural framework for understanding its activation and selective autoinhibition.

## Introduction

*Pseudomonas aeruginosa is a* versatile opportunistic pathogen, capable of causing serious infections, many of which are associated with biofilm formation. Extracellular proteases and peptidases are prominent components of *P. aeruginosa* biofilms^1^ and are also produced during chronic airway infections^2–4^, where they contribute to bacterial physiology and pathogenesis. Secreted lytic virulence factors facilitate the establishment of infection and host damage, particularly in wounds and burns^2,5–7^ and secreted proteases are considered major components of this virulence repertoire. In cystic fibrosis-adapted strains associated with chronic infections, proteolytic activity can enhance pathogenicity by weakening immune defences and promoting tissue invasion^8^. Moreover, bacteria belonging to biofilm communities surviving antibiotic treatment, retain high levels of protease secretion^9^.

Many secreted proteases are produced as inactive zymogens, which require proteolytic processing to become fully functional enzymes. In *P. aeruginosa,* elastase B (LasB), the lysine-specific endopeptidase (locus tag PA14_09900, also known as PrpL, LysC and Protease IV or PIV, and hereafter referred to as PIV) and the aminopeptidase PaAP are thought to act in tandem in an activation cascade that ultimately results in mature forms for these enzymes^10^. This places PIV within a broader secreted protease network in which proteolytic processing regulates the activity of other extracellular enzymes.

Lysine-specific endopeptidases such as PIV cleave the peptide bond immediately after lysine residues (Lys-Xaa)^11^. This stringent specificity has resulted in these proteases being highly valuable tools used for biotechnology and mass spectrometry applications^12^. PIV is a highly conserved extracellular trypsin-like serine protease that is widely distributed among *P. aeruginosa strains*^13^. It contributes to virulence in multiple host systems, including plants and mammalian infection models, and is a well-established virulence determinant in corneal infections^14,15^ where it promotes tissue damage and bacterial pathogenicity.

The PIV gene (*prpl)* encodes a 48 kDa zymogen containing an N-terminal signal peptide (Pre-Pro-PIV). The signal peptide is required for extracellular secretion, and its removal produces an approximately 45 kDa proenzyme species (Pro-PIV), which is subsequently processed extracellularly to release the fully active, ∼27 kDa mature protease. Although removal of the propeptide region is widely accepted as a prerequisite for PIV maturation, the route by which this occurs remains unresolved^16^. Studies using different combinations of deletion strains have implicated extracellular proteases including LasB in pro-PIV processing^10,17–19^ and the isolated PIV propeptide has been reported to inhibit the mature enzyme in trans^17^. Conversely, active 27 kDa mature PIV was isolated from strain PA103-29, which lacks LasB production^11^, and catalytic-site substitutions prevented PIV autoprocessing^20^, which are both observations consistent with an intrinsic auto-activation pathway. To our knowledge, purified recombinant mature PIV has not previously been reported, limiting detailed investigation of its structure, mechanism and self-activation.

Despite the important role of PIV as a well characterised virulence factor, the structural basis of its precursor inhibition and the sequence of events leading to maturation remain poorly understood. Here we determined a high-resolution crystal structure of the inactive PIV precursor and identify a previously uncharacterised Cap–CUB pro-region that occludes the catalytic cleft. Using recombinant expression in a host lacking the relevant *Pseudomonas* extracellular proteases, we establish the production of mature active PIV and examine its sequential self-processing. We combine these structural observations with kinetic assays using Cap-derived peptides as well as established serine-protease inhibitors, and assess the capacity of mature PIV to activate PaAP. Together, these data provide a framework for understanding PIV autoinhibition, extracellular maturation and substrate recognition, and establish a foundation for future structure-guided design of inhibitors targeting this important *P. aeruginosa* virulence factor.

## Results

### PIV adopts a three-part autoinhibited architecture

To define the organisation of the PIV precursor, we crystallised a protein construct lacking the signal peptide and carrying the catalytic substitution S409A. The structure was determined at 2.17 Å resolution in space group C 2 2 2_1_, with four PIV molecules in the asymmetric unit. NanoDSF and DLS measurements indicated that the protein is monomeric in solution (Figure S1). Electron density was sufficiently well defined to model residues 29– 462 in each chain, with the exception of a poorly ordered region around residues 70-80. The four crystallographically independent molecules superpose closely (root-mean-square deviation (RMSD) <0.5 Å across approximately 420 Cα atoms).

The precursor comprises three contiguous structural regions (Figure 1A&B): an N-terminal Cap region (residues 25-90), a CUB domain (PFAM: PF00431, residues 91–211) and a C-terminal protease domain (residues 212–462). The protease domain forms the two-β-barrel architecture characteristic of trypsin-like S1-family serine proteases. Above it, the CUB domain forms a compact, predominantly β-sheet-rich module. The Cap region is less globular. It extends from the N terminus to wrap around the whole structure like a clamp traversing the catalytic cleft before reconnecting with the CUB core. In this arrangement, the CUB domain provides a scaffold while the extended Cap region acts as a tethered clamp across the enzyme surface.

**Figure 1.**
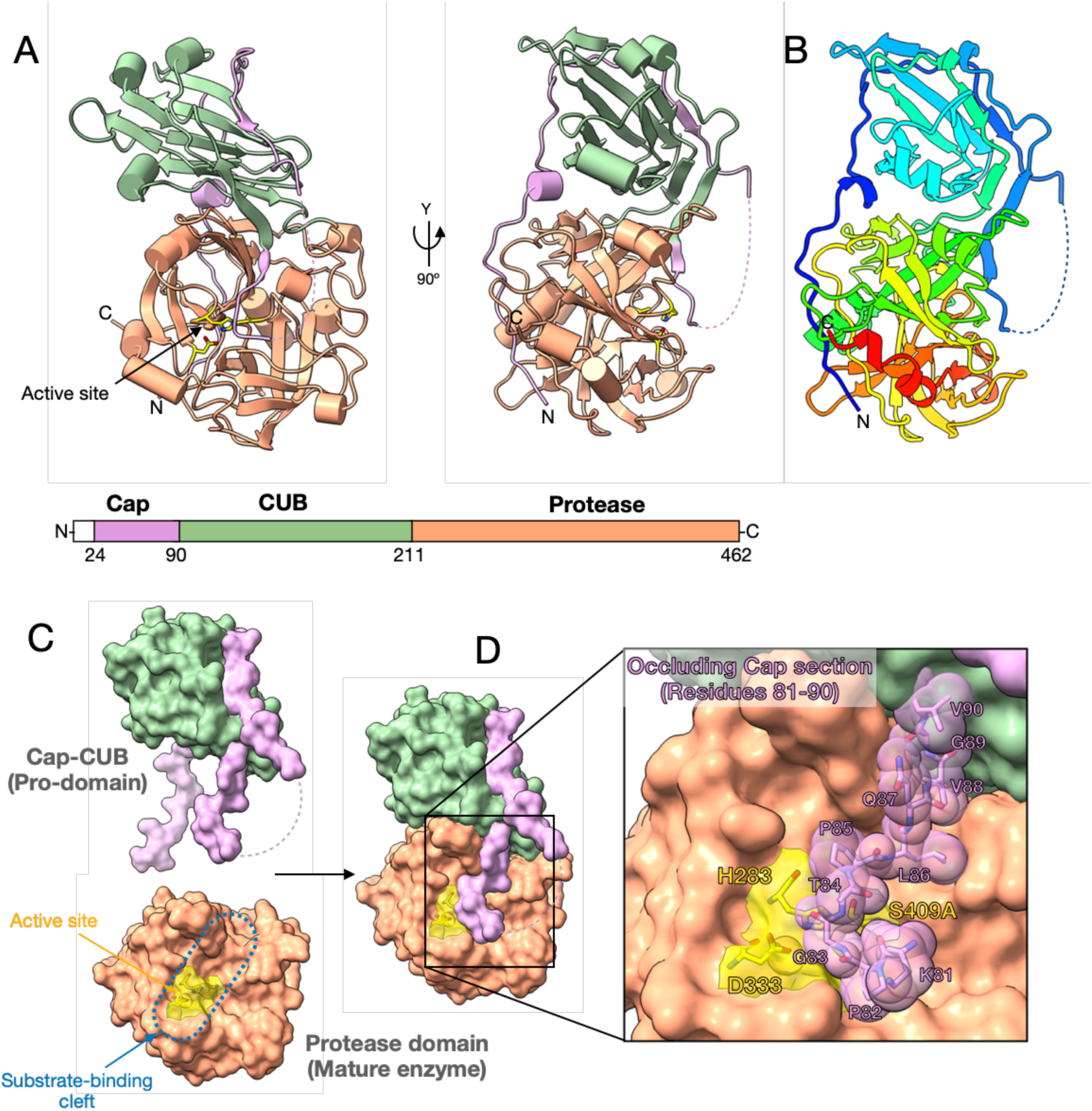
Structure of PIV reveals the autoinhibitory mechanism. **(A)** Orthogonal ribbon views of PIV_S409A_, coloured by domains: Protease domain (tan), the CUB domain (green) and the Cap domain (plum). Both Cap and CUB domains make up the pro-peptide which is released during the maturation process, to yield the active protease domain. The structure reveals the self-inhibitory mechanism of the pro-peptide, whereby the active site is occluded by the Cap domain. The Cap domain wraps around the protease and CUB domains acting like a molecular tie. This binding mode blocks access to the active site and acts to keep the enzyme in an inactive state. **(B)** Ribon diagram coloured in rainbow from N-terminus (Blue) to C-terminus (Red) showing the continuity of the fold. **(C)** Linear schematic showing the domain organisation of PIV, including the signal peptide, pro-domains (Cap & CUB domains) and catalytic protease domain. Residue numbers indicate the domain boundaries. Colours correspond to those used in the three-dimensional structure throughout the manuscript. **(D)** Structure of PIV, broken down into Cap-CUB (pro-domains) and Protease (mature enzyme) components, highlighting how these components interact to block the catalytic site. The structures are in surface view and coloured according to domains. Active site residues are highlighted in yellow (through transparent surface) and the substrate-binding cleft is marked by dashed blue line. **(E)** PIV structure in surface view, coloured according to domains. The full-length structure highlights how the Cap domain blocks access to the preformed active site by occupying the substrate-binding cleft. To the right is a zoomed in view of this interaction. Residues 81-90 of the Cap domain are displayed and the catalytic residues are highlighted in yellow. The proline rich region of Cap sits above the substrate-binding cleft and blocks access to the active site.

The domain arrangement provides a molecular rationale for the inactivity of the full-length precursor PIV. Surface views show that residues 81–90, at the Cap–CUB junction, lie directly above the catalytic residues and physically block the substrate-binding cleft (Figure 1C&D). The Cap therefore does not disrupt formation of the protease fold; instead, it masks a pre-organised catalytic surface. The independently determined 3.30 Å structure of mature PIV in complex with its separated propeptide (PDB 9OMD) adopts the same overall arrangement^19^. Superposition gives RMSDs of 1.68 Å across 250 residues of the protease domain and 2.06 Å across 169 residues of the pro-region, supporting a conserved precursor conformation. The present higher-resolution data, resolves the active-site and interdomain contacts in greater detail, providing structural information of the pre-activated & unprocessed form.

### PIV auto-inhibitory mechanism

Structural information is available for the mature forms of homologous lysyl endopeptidases, enabling a comparison and description of unique features of PIV. High resolution structures of *Achromobacter lyticus* (PDB 4GPG) and L*ysobacter enzymogenes,* (PDB 1ARC) are solved in complex with the covalent inhibitor TLCK, which binds at the catalytic site. These structures revealed important features of the catalytic mechanism and also substrate selection. Comparing our auto-inhibited structure to these mature enzyme structures, informs on the structural basis for how the pro-peptide region blocks access to the preformed catalytic site.

Our structure was obtained by mutating the catalytic serine residue to an alanine, which results in a full-length protein, incapable of self-processing. Without this mutation, during protein expression and subsequent purification, the enzyme self-processed to a ∼26 kDa species (discussed below). Similarly, mutations of the other two residues forming the catalytic triad (D333 and H283), also resulted in full-length and unprocessed protein (Figure S2). Activity is regulated by the presence of the pro-domain, which occludes the active site, with the Cap domain wrapped around the whole protein. While the Cap domain is largely unstructured, the CUB domain has a conserved fold, present in many peptidases belonging to MEROPS families M12A (astacin) and S1A (chymotrypsin).

The PIV active site lies in a shallow, elongated cleft at the interface between the two β-barrels of the protease domain (Figure 1A). It contains the catalytic triad H283, D333 and S409; the nucleophile is represented by alanine in the crystallised S409A variant. Importantly, the remaining active-site geometry is retained in the precursor, indicating that inactivity arises principally from restricted substrate access rather than gross distortion of the catalytic domain.

To examine how a lysine-containing substrate would engage this surface, we modelled the peptide AAAKAAA with the PIV protease domain using AlphaFold 3^21,22^. The model places the P1 lysine side chain in the narrow, electronegative S1 pocket, where its terminal ε-amino group is positioned to interact with polar groups including S223 and D234 (Figure 2A&D). The peptide backbone spans the surface groove, and the scissile-bond carbonyl is directed towards the main-chain amide donors that form the oxyanion hole (Figure 2D). This pose is consistent with the established preference of lysyl endopeptidases for lysine over the longer guanidinium-terminated side chain of arginine. Because this is a predicted complex rather than an experimentally determined substrate-bound structure, care must be taken to avoid overinterpretation of binding contributions from distinct groups. Therefore, it proposes a plausible binding complex rather than atomic interaction energies.

**Figure 2.**
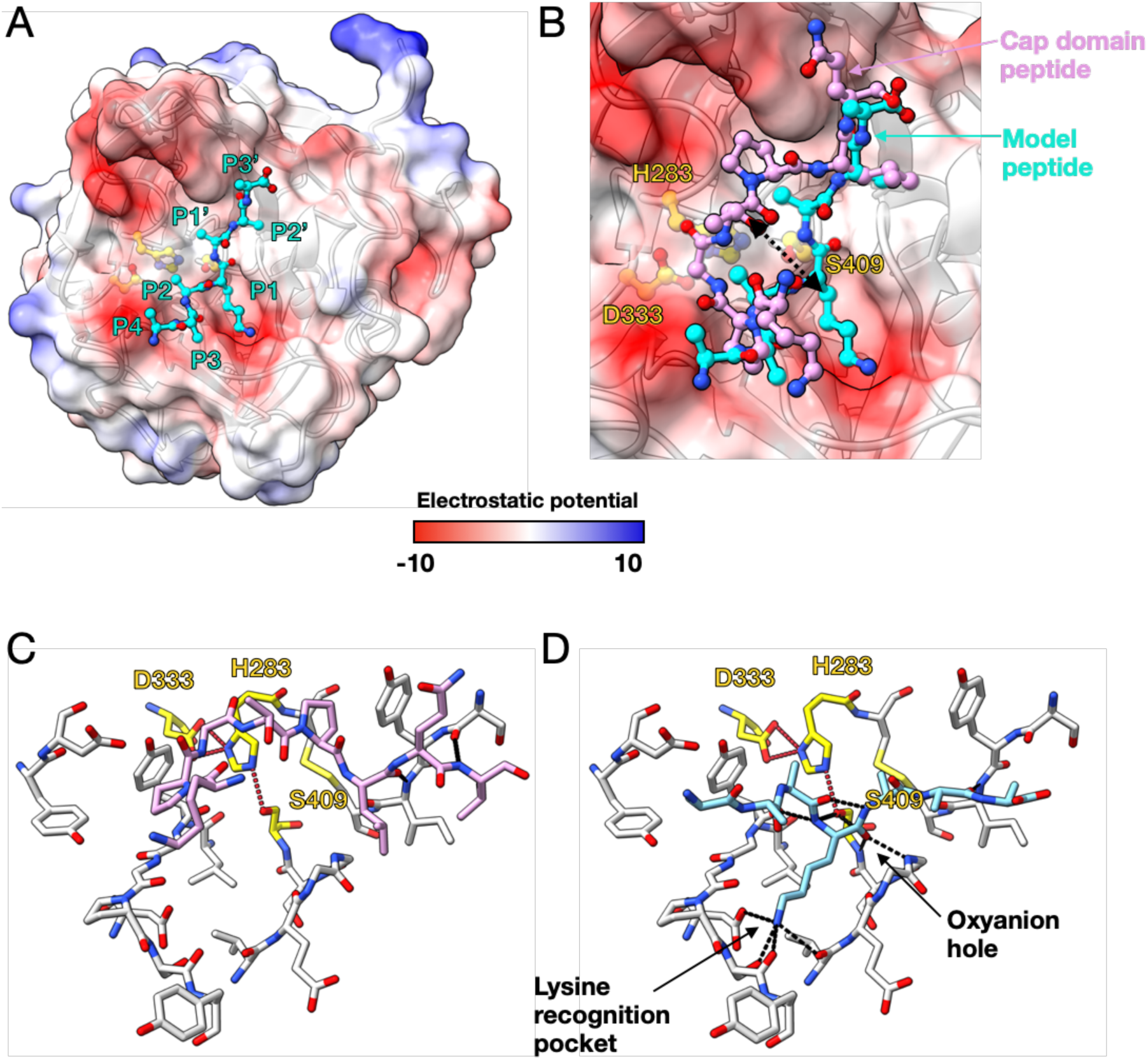
Substrate binding site and occlusion by the cap domain. **(A)** Structure of PIV protease domain only in surface representation, coloured according to electrostatic potential. Binding of a peptide (AAAKAAA) was modelled to PIV using AlphaFold3. The modelled peptide (cyan) is modelled to bind similarly to other trypsin-like enzymes, into the shallow cleft running along the surface. The substrate specificity residue (Lys, P1) binds in the S1 pocket and the side chain amide group interacts with an electronegative pocket. **(B)** A close of view of the interaction between PIV and the modelled substrate (cyan) in comparison to the binding mode of the Cap domain (plum). The Cap domain does not sit in the cleft but sits atop of the cleft. Two Proline residues facilitate this binding mode. **(C)** The PIV substrate binding pocket residues are shown in stick form. Catalytic residues are shown in yellow with hydrogen bonds shown by red dashed lines. The Cap domain is not positioned in the substrate binding cleft and makes few hydrogen bonds (black dashed lines) with the substrate binding pocket residues. **(D)** In comparison, the AlphaFold3 modelled substrate (cyan) binds into the cleft in a catalytically competent position. The modelled substrate is predicted to participate in several hydrogen bonds (black dashed lines) with the substrate binding site residues. The oxyanion hole and the lysine recognition pocket are highlighted.

Superposition of the modelled peptide with the intact precursor reveals two distinct binding modes (Figure 2B). The peptide descends into the catalytic groove, occupies the S1 pocket and approaches the oxyanion hole. By contrast, residues 81-90 from the Cap region remain above the groove and make comparatively few direct contacts with its specificity pockets. It therefore acts as a steric lid, not as a conventional pseudosubstrate. The Cap cannot be accommodated simultaneously with the modelled peptide and would prevent a polypeptide substrate from adopting a catalytically competent trajectory.

The structure can be visualised as having 2 buried surface areas, one between the protease and CUB domain and another between the Cap and the protease-CUB complex. The buried surface area was calculated use PBDePISA^23^ which shows surface areas of 2545 Å^2^, between the Cap and Pro-PIV and a buried surface area of 931 Å^2^, between the CUB and PIV domains.

Structurally, the Cap and CUB domains possess distinct structural features, and structure-based searches with DALI^24^ and Foldseek^25^ identified divergent CUB-containing proteins but no close match to the complete Cap–CUB–protease arrangement. Thus, although the CUB fold is not unique to PIV, its integration with an extended active-site-blocking segment appears to define a specialised regulatory module. Whether the CUB surface also contributes to substrate recruitment or localisation before proteolytic release remains to be tested. The CUB domain is an unusual structural feature, as it does not directly occlude the active site but its combination with the Cap acts as a “helmet with straps”, with the straps blocking the active site. CUB domains are associated with several proteins and facilitate protein:protein and protein:ligand interactions. Searches for structural matches of this domain reveal several divergent proteins, including 1SPP, in which the CUB domain is required to recognise receptors, and functionally unrelated sugar binding proteins. Together, these suggest the CUB domain of PIV may assist in localisation of the protease to its substrates, before activation. Our analysis shows that the enzyme is only fully active when cleavage to a 26 kDa species occurs (i.e. removal of cap and CUB pro domain).

### PIV undergoes auto-lysis leading to protein maturation

Substrate profiling of different lysyl endopeptidases homologues exploring their deployment in proteomic workflows revealed strict preference for lysine for PIV, with under 0.4% cleavage taking place following other amino acids^26^. Compared to homologues from *Achromobacter lyticus*, and *Lysobacter enzymogenes*, PIV showed a bias against proline, glutamic acid, and additional lysine residues on positions following the cleavage site (P1), indicating a potential preference to some flanking residues in detriment of others. During initial purification attempts, we observed extensive self-cleavage of PIV, with the generation of a “3-protein band” intermediate that continues to presumably self-cleave and give rise to lower molecular weight contaminants during purification (Figure S3). Figure S4 depicts all lysine residues present in PIV as well as their flanking sequences.

To generate highly pure and recombinantly expressed PIV, we developed a double affinity protocol employing a Streptactin tag on the protein’s N-terminus as well as a his-tag in the protein C-terminus. However, alone this was not sufficient to overcome issues in the separation of mature and intermediate enzyme forms during purification. We hypothesised that correct protonation of residues in the catalytic triad was essential for enzyme function during expression and purification, as PIV does not possess carboxylates neighbouring the catalytic serine, which are known to enable catalysis at low pH in Cathepsin A and other serine proteases^27^. Therefore, we envisioned a pH-controlled strategy to trigger self-cleavage in a controlled manner. Purification of Strep and his-tagged PIV at low pH (6.5) enabled the generation of a stable “3-protein band” population (Figure 3B and Figure S3). Incubation at room temperature for 3 days either at pH 5.8 or 8.5 was required to generate fully matured PIV (Figure 3A). Mass spectrometry of intermediate bands allows the proposal of a sequential cleavage sequence that ultimately leads to fully mature PIV. Initial steps in the activation are slow (Figure 3B), with full-length PIV possessing a half-life of 2.3h, leading to a stationary level of the a “3-protein band” intermediate state, which then underwent rapid processing on day 3. Additional data on mass spectrometry of PIV variants and activation intermediates is available on Figure S5.

**Figure 3.**
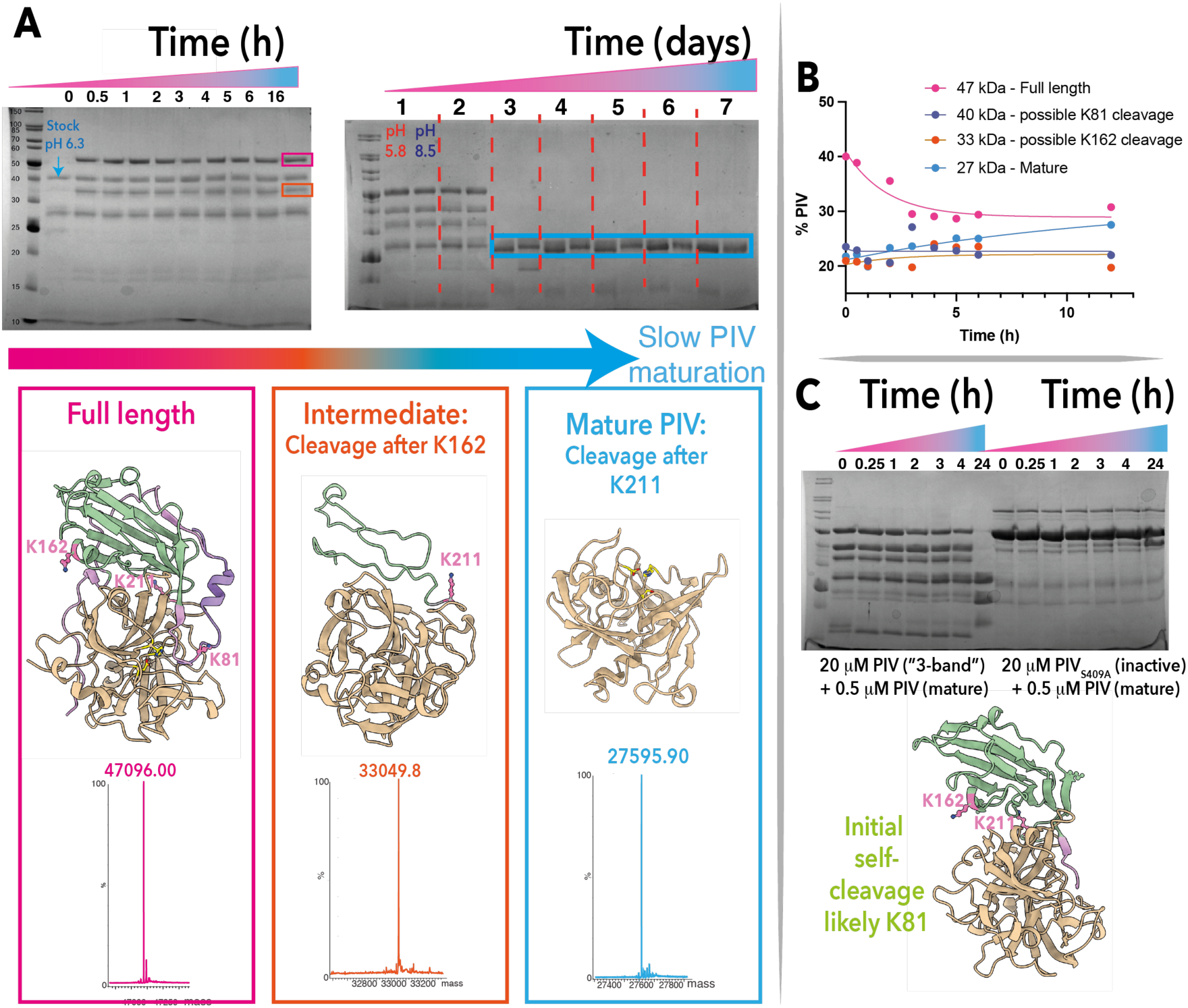
PIV can self-activate. For all structure cartoons showed, the N-terminal inhibitory cap is shown in plum, CUB domain in green, and catalytic domain corresponding to mature PIV in tan. Residues 70-80 could not be modelled, and an overlaid Alphafold3 prediction for this short region is shown in purple. **A**) SDS-PAGE depicting a time course for PIV self-activation at pH 8.5 (shorter time spans of hours), or pH 5.8 and 8.5 for longer time spans (days). At indicated times protein solution was also analysed by mass spectrometry, enabling the identification of full-length PIV (pink box, observed 47096 Da, calculated 47098.42 - Δmass = -2.3 Da), an intermediate band likely following cleavage at K162 (orange box, observed 33049.8 Da, calculated 33057.4 - Δmass = 7.6 Da), and mature PIV after cleavage on K211 (blue box, observed 27595.9 Da, calculated 27598.2 - Δmass = -2.3 Da). Possible cleavage sites are listed on Figure S4. Structure for full-length PIV is from this work, while intermediate and mature PIV were generated with Alphafold3. Both incubation of the protein at room temperature at low (5.8) or high pH (8.5) led to complete activation on day 3. **B**) Quantification of band intensity in function of time from the SDS-PAGE shown in panel A. A clear decay of the band corresponding to the full-length PIV is observed, with slow formation of intermediates leading to mature (fully active) PIV, corresponding to cleavage after K211. **C**) When active site mutant PIV_S409A_ (catalytically inactive) is used as a substrate for mature PIV, no significant activation is observed, pointing to a first intra-molecular cleavage, which could take place on K81 as it is closest to the active site. For SDS-PAGE on panels A and C, the first lane is the molecular weight marker (Thermo Scientific PageRuler™ Unstained).

To test whether cleavage took place in trans, we used inactive mutant PIV_S409A_ or “3-protein band” intermediates as substrates for mature PIV (Figure 3C). After 24h the “3-protein band” intermediates were converted into fully matured PIV, while no or little visible cleavage took place for PIV_S409A_. Our crystal structure shows a flexible linker between the Cap and CUB domains, in which density for K81 is unresolved. Therefore, cleavage data are consistent with an activation step that is triggered by an initial cleavage in cis. We propose this is likely on K81 as other lysine residues are distant from the active site and in well-structured regions.

### PIV is covalently inhibited by TLCK, and weakly inhibited by Cap peptides

Other work employed PMSF as a broad serine protease inhibitor to inactivate PIV^28^ but an inhibition experiment with excess PMSF did not show detectable inhibition (Figure 4A), and no covalent modification was observed when PMSF was added during its purification (Figure S5).

**Fig. 4:**
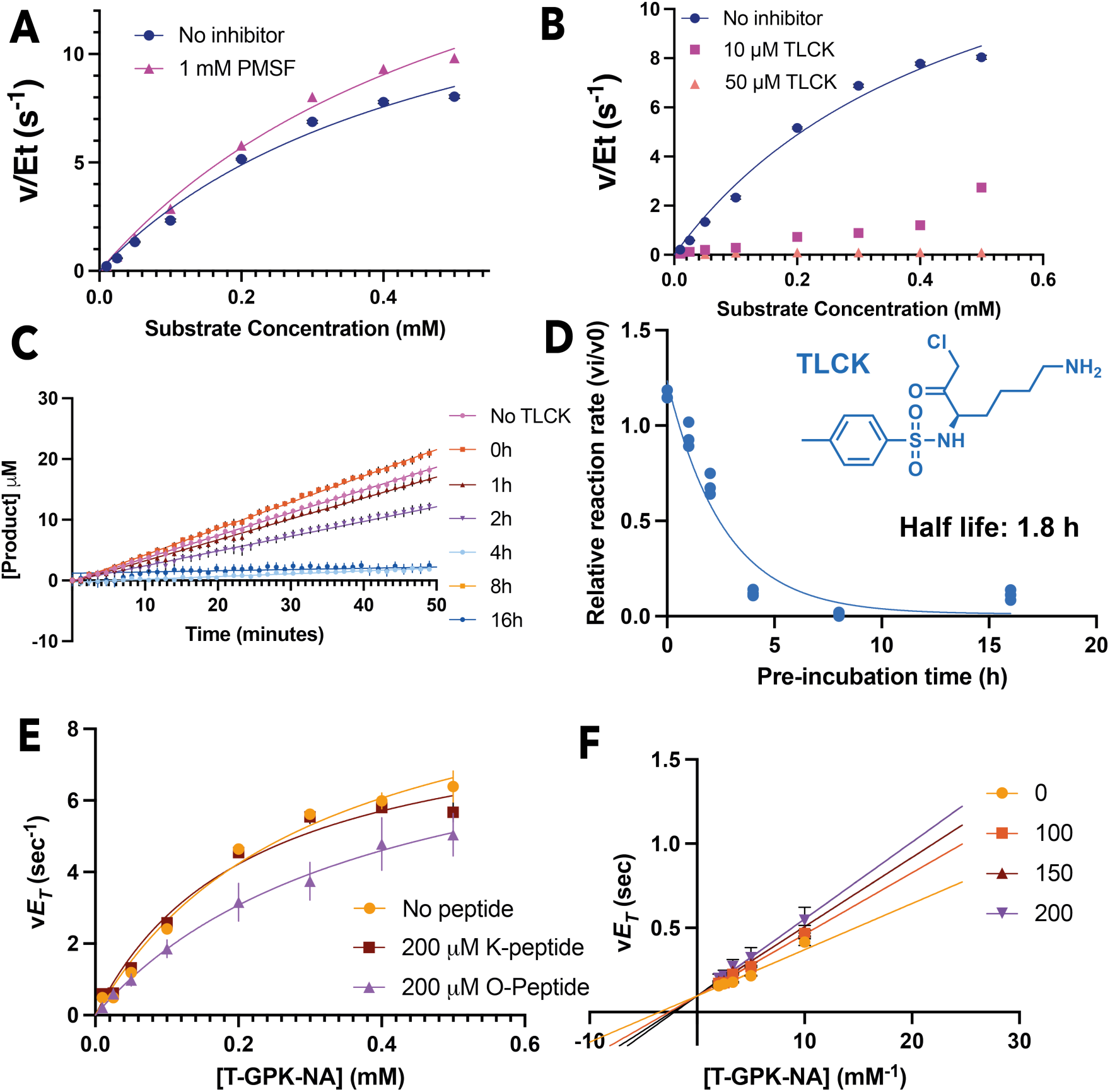
PIV inhibition. **A)** PMSF is not an inhibitor for PIV under the conditions tested. **B)** TLCK is an inhibitor for PIV, with complete inhibition at a higher concentration. **C)** Pre-incubation of PIV and TLCK for different amounts of time, followed by inhibitor jump dilution shows that **D)** residual activity is reduced in function of pre-incubation time. Fit is to an exponential equation, to yield an observed inhibition rate of 3.8 h^-1^, and a half-life of 1.8 h. **E)** A peptide sequence corresponding to the Cap region is not an inhibitor for PIV (K-peptide), while the same sequence carrying an ornithine residue instead of a lysine is a weak inhibitor (O-Peptide). **F)** O-peptide is a competitive inhibitor, with an approximate *K*_i_ value of 300 μM. Lines are fitted values to a competitive inhibition equation. Non-linearized data are present on Figure S7.

TLCK is a well-established serine protease inhibitor for trypsin and related proteases which covalently modifies the catalytic histidine residue, ultimately leading to enzyme inactivation. TLCK demonstrated incomplete PIV inhibition at 10μM, but complete inhibition at 50 μM (Figure 4B). To confirm covalent inhibition of PIV, we carried out a jump dilution experiment pre-incubating enzyme with excess TLCK for different amounts of time, followed by rapid dilution into a standard reaction mixture with excess chromogenic substrate T-GPK-NA (Nα)-tosyl-glycyl-prolyl-lysyl-4-nitroanilide – Figure 4C). Residual activity reduced as pre-incubation time increased, characteristic of a slow covalent inactivation step, with a half-life of 1.8 h when TLCK is present at a 2:1 ratio over PIV (Figure 4D). Increasing the inhibitor excess during incubation to a 20:1 ratio let to complete inactivation after 4h (Figure S6a).

We confirmed covalent modification by intact protein mass spectrometry (Figure S6b) as overnight incubation with TLCK led to the observation of a protein species with additional 297 Da, consistent with TLCK covalent modification of PIV.

Because the structure of PIV revealed the Cap sequence to cover the enzyme’s active site, we tested whether peptides harbouring the Cap sequence were inhibitors. Peptides lacking a lysine residue were not inhibitors, while the inclusion of an ornithine residue led to weak inhibition by these peptides, via a competitive mechanism towards the substrate (Figure 4e and f). Ornithine-containing peptides showed inhibition at the high micromolar range (*K*_i_ estimated to be 300 μM), but could not be cleaved by PIV, while the same peptide sequence harbouring a lysine was efficiently cleaved under the same conditions (Figure S8).

### PIV can activate the aminopeptidase PaAP

Previous work implicated PIV in an activation cascade to remove an inhibitory C-terminal sequence from PaAP in *P. aeruginosa*^29^. Since PIV cannot cleave C-terminal to residues other than lysine, it does not catalyse the hydrolysis of the PaAP substrate Leu-pNA. When active PIV is added to an essay mixture including PaAP and Leu-pNA, there is an increase in the apparent rate of the reaction as the concentration of PIV increases, ultimately reaching a plateau (Figure 5). This could be due to a rate limiting process in the activation cascade which is independent of the concentration of PIV. This result demonstrates that it can perform this activity and ultimately increase the activity of PaAP. Proteomic analysis of secreted proteins from *P. aeruginosa* (Figure S9)^30^ in cells growing in culture shows the last 24 residues from the C-terminal tail of PaAP to be absent in its mature form.

**Figure 5:**
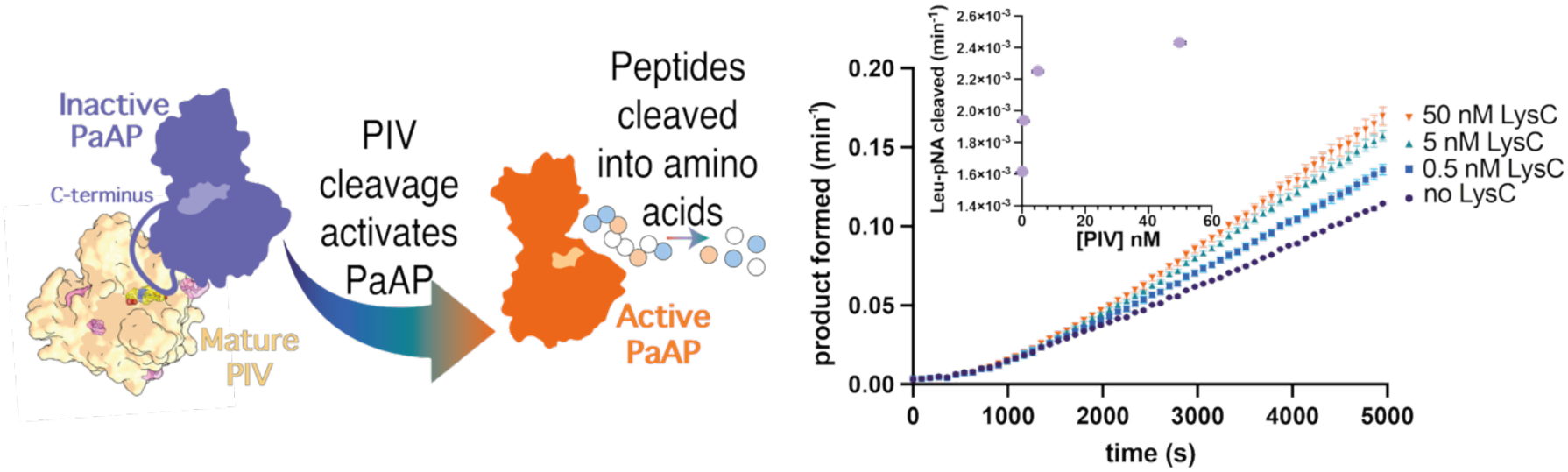
PIV can cleave PaAP to increase the activity of the aminopeptidase. Left, proposal for PIV cleavage of PaAP C-terminal tail. Right, assay with PaAP in the presence of PIV, showing a concentration dependent increase in the rate of cleavage of the aminopeptidase substrate Leu-pNA. The inset shows a plateau on the rate increase, likely due to other factors contributing to activation, so that that generation of active PaAP is limited not only by the encounter of PIV-PaAP.

## Discussion

Here, we present the structure of pro-PIV, revealing the structural mechanism of precursor regulation within the established family of lysine-specific, trypsin-like serine proteases. The catalytic protease domain of PIV retains the two-β-barrel fold, catalytic triad and lysine-selective S1 pocket found in related homologues, *Achromobacter lyticus* protease I (PDB 1ARB, 1ARC and 4GPG)^31,32^ and *Lysobacter enzymogenes* LysC (PDB 4NSY and 4NSV)^33^. However, these structures lack the N-terminal regulatory regions present in PIV (31RM). These mature structures represent the exposed, substrate-accessible state of the enzyme family. By comparison, the PIV precursor retains the pre-organised catalytic site, which is sterically blocked by the N-terminal Cap–CUB pro-domain assembly. Consequently, the primary mechanism of PIV autoinhibition is steric occlusion of a conserved substrate-binding surface, rather than reorganisation or reconstruction of the catalytic site.

The overall architecture is consistent with the recently reported 3.30 Å PIV structure (PDB 9OMD), supporting a conserved association between the pro-region and protease domain. However, 9OMD structure was assembled from separately purified pro-region and protease components, whereas our 2.17 Å structure captures the Cap, CUB and protease regions as a covalently continuous precursor, thus representing an unprocessed form. The higher-resolution data provides more confident definition of the catalytic centre, substrate-binding cleft and interdomain contacts. Together, these features show that the CUB domain provides a structural scaffold for an extended Cap that crosses above the catalytic cleft. The Cap does not occupy the specificity pockets in the manner of a conventional pseudosubstrate; instead, it forms a tethered steric barrier that prevents a polypeptide substrate from adopting a catalytically competent trajectory. These two PIV structures complement each other, representing pre- and post-cleaved states. The main region of differences arise from residues 206-217 (including the final cleavage site of K211) and residues 65-81 (including the disordered loop region of the cap domain).

Together, the structure and processing data support a sequential mechanism of PIV activation. Wild-type PIV underwent time-dependent self-processing through discrete intermediates, whereas substitutions within the catalytic triad retained the full-length precursor. Mass spectrometry supports cleavage after K162 and a final cleavage after K211, generating the ∼27.6 kDa mature enzyme. Proteomic analysis of secreted proteins provides in vivo corroboration that cleavage at K211 results in mature PIV, as this is the species that is identified, and peptides corresponding to the Cap and CUB domains were not detected (Figure S9). The initial activation step appears mechanistically distinct and far slower than subsequent processing steps. Mature PIV readily converted intermediate species into mature enzymes in trans. However, mature enzyme showed little activity towards the intact PIV_S409A_ precursor. Initiation may therefore require an unfavourable conformational change or state that permits intramolecular cleavage. K81 is a plausible candidate as it lies close to the catalytic cleft within the poorly ordered Cap–CUB linker, although cleavage at this site has not been demonstrated directly. Such an event could weaken the attachment of the Cap and expose downstream sites, allowing faster intermolecular cleavage after K162 and K211.

Neighbouring residues in different putative cleavage sites within PIV provide a possible explanation for these different rates. Mass-spectrometric profiling of lysine-specific peptidases, including PIV, identified sequence features associated with efficient cleavage or missed cleavages^26^. The sequence surrounding K81 (PLAP**K**↓PGTP), contains Pro residues at P1′ and P2, which may restrict the conformation required for productive substrate binding^26^. Mapping these preferences onto the accessible lysines in PIV similarly identifies relatively unfavourable environments around K81 and K93, whereas K154, K162 and K211 are surrounded by residues associated with more efficient processing (Figure S10). The sequence surrounding the proposed initiating site may therefore impose an additional barrier. Once this barrier is overcome and the Cap–CUB assembly is destabilised, the more favourable downstream sites would become accessible for rapid processing in trans. Environmental conditions or extracellular proteases such as LasB could accelerate activation by facilitating or bypassing this slow initiating step.

The behaviour of PIV during purification is also consistent with a kinetic barrier rather than an absolute block to activation. Early preparations underwent extensive self-cleavage (Figure S3), whereas purification at pH 6.5 stabilised a partially processed population that matured slowly during subsequent incubation. Although trypsin-like serine proteases commonly favour neutral-to-alkaline conditions^34,35^, PIV ultimately matured at both pH 5.8 and pH 8.5. Although mechanistically distinct from PIV, pH-dependent conformational trapping and slow recovery have been described in serpin systems^36^, and a pH-denatured subtilisin variant can slowly promote intermolecular maturation of an inactive precursor after transfer to neutral pH^37^. Thus, pH influences the rate at which the precursor escapes autoinhibition but does not operate as a simple on–off switch. The prolonged lag, followed by more rapid loss of intermediates, is instead consistent with conformational sampling of a poorly accessible initiating site followed by propagation through more favourable cleavage sites.

Auto-processing and the steps involved in the activation of PIV has been widely discussed and contested^16^. However, our results support a sequential auto-processing model, previously suggested when mature PIV was isolated from PA103-29 which lacks LasB production^11^ and catalytically-inactivated enzyme failed to mature^20^. The recombinantly expressed system used here, removes the complexity and ambiguity surrounding activation steps, by removing other complicating extracellular protease activity.

Together, our findings strongly support self- or auto activating not mandatorily reliant on other extracellular enzymes, e.g. LasB^10,17–19^, although cleavage via extracellular proteases within the disordered Cap region may accelerate processing. LasB and/or AprA activity may bypass the kinetically slow initiating step, whereas self-processing provides an autonomous route to maturation when these enzymes are absent or limiting. PIV activation may therefore be controlled through alternative but convergent pathways, all of which ultimately remove the Cap–CUB assembly and expose the mature lysyl-endopeptidase domain.

This mechanism has broader implications for the extracellular protease network of *P. aeruginosa*. Mature PIV contributes to virulence through the degradation of host structural and immune proteins, but it can also regulate other secreted bacterial enzymes. PaAP is a biofilm-associated aminopeptidase whose activation requires removal of an inhibitory C-terminal sequence^29^, previously proposed to occur due to PIV cleavage at K512–A513^10^ and confirmed here (Figure 5, Figure S9). The concentration-dependent increase in PaAP activity observed here is consistent with this processing pathway and positions PIV as an intermediate component of an extracellular proteolytic cascade. Activation of a relatively small population of PIV molecules could therefore amplify proteolytic activity through subsequent maturation of PaAP and potentially other secreted substrates.

Together, these findings establish PIV is secreted as a precursor protein which contains a mature-like catalytic domain whose substrate-binding cleft is sterically occluded by a pro-region, composed of a Cap–CUB domain assembly. Sequential processing removes this inhibitory assembly through kinetically distinct initiation and propagation phases, producing the stable mature enzyme. The structural basis of autoinhibition, together with incapacity to cleave ornithine-substituted peptides, provides a framework for developing non-cleavable substrate mimetics that engage the lysine-recognition pocket. Such mechanism-guided inhibitors could selectively disrupt PIV-dependent proteolytic cascades and offer a route to attenuating *P. aeruginosa* virulence.

## Methods

### Materials, chemicals and general

Chemicals used in this investigation were purchased from Fisher and Merck. Peptides were purchased from Genscript at >98% purity and used without further purification. Primers used for mutagenesis were purchased from IDT. A list of primers used in this study is provided in Table S1 and a list of plasmids is provided in Table S2. Kinetic data were fitted using GraphPad Prism. SDS-PAGE were imaged using a Biorad Chemidoc XRS+ and analysed by ImageJ.

### Strains, media, and growth conditions

Strains used in this study are listed in Table S3. *E. coli* strains were grown in LB media, supplemented with appropriate antibiotic, at 37 °C.

### Cloning, expression and purification of PIV

For the purification of PIV inactive mutants, a codon optimized sequence encoding the secreted form of PIV (comprising residues 25-462 of Uniprot entry: Q02SZ7 from *Pseudomonas aeruginosa* PA14) was used, lacking the predicted N-terminal signal peptide region (residues 1-24). This gene construct was cloned into pJ411, in line with an ATG start codon and immediately followed by a C-terminal His_8_ affinity purification tag (EHHHHHHHH). Mutant PaPIV_S409A_ was generated using a using an in-house site-directed mutagenesis protocol adapted from the Q5^®^ Site-Directed Mutagenesis Kit (New England Biolabs, E0554). Constructs and mutations were confirmed by Sanger sequencing using Eurofin Genomics before transformation into the *E. coli* expression strains.

pJ411::PaPIV_S409A_ constructs were transformed into SHuffle® T7 by NEB, to assist disulphide bond formation. Transformed cells were grown at 37 °C (shaken at 180 rpm) in LB media (10 g/L NaCl, 10 g/L tryptone, and 5 g/L yeast extract) supplemented with 100 μg/ml Kanamycin until an OD_600_ of 0.8 was reached. Gene expression was induced with 1 mM IPTG and incubated over night at 16 °C (shaken at 180 rpm). Cells were harvested by centrifugation at 8855 g (Beckman JLA 8.1000 rotor) for 10 min and the pellets stored at −20 °C.

Cells were re-suspended in fresh lysis buffer (50 mM HEPES pH 8.0, 20 mM Imidazole pH 8.0, 300 mM NaCl) and incubated with ∼1 mg/ml lysozyme for 30 min at 4 °C. Cells were lysed using a high-pressured cell disruptor. Insoluble cell debris was removed by centrifugation for 30 minutes at 33,000 g (Beckman JA 25.50). The supernatant was loaded onto a 5 ml HisTrap HP column (GE Healthcare) pre-equilibrated with lysis buffer. The column was washed with 20 column volumes (CV) of lysis buffer to remove non-specific interacting proteins. 5 CV of elution buffer (50 mM HEPES pH 8.0, 250 mM Imidazole pH 8.0, 300 mM NaCl) was passed over the column to elute the His_8_-tagged PIV. Fractions containing pure PIV, evaluated by SDS-PAGE, were pooled and dialysed over night against 2 L of dialysis buffer (25 mM HEPES pH 8.0, 50 mM NaCl, 50 mM KCl). Pooled & dialysed samples were concentrated using 30 KDa MWCO spin concentrators to ∼10 mg/ml for further experiments. Protein concentration was deduced from A280 readings using a Denovix DS series instrument.

For production of the active PIV, a strep tag (MG**WSHPQFEK**G, sequence in bold) was added to the N-terminus of the pJ411::PaPIV construct using the same in-house site-directed mutagenesis protocol. This generated a protein with an N-terminal Strep tag and a C-terminal Histag. Constructs were transformed into *E. coli* BL21. Plasmid DNA was extracted using the Promega PureYield™ Plasmid Miniprep system and sequence and identity confirmed by Oxford Nanopore sequencing (MicrobeNG plasmidseq).

Cells were transformed with pJ411::PaPIV plasmid and grown at 37 °C (shaken at 180 rpm) in LB media (10 g/L NaCl, 10 g/L tryptone, and 5 g/L yeast extract) supplemented with 100 μg/mL Kanamycin until an OD600 of 0.5-0.8 was reached. Gene expression was induced with 0.75 mM IPTG and incubated over night at 18 °C (shaken at 180 rpm). Cells were harvested by centrifugation at 8855 g (Beckman JLA 8.1000 rotor) for 20 min and the pellets stored at −20 °C.

Cells were resuspended in lysis buffer (50 mM MES pH 6.5, 20 mM Imidazole, 300 mM NaCl) and incubated with 1 mg mL^-1^ lysozyme for 60 min at 4 °C. Cells were lysed using a high-pressured cell disruptor (25 kpsi). Insoluble cell debris was removed by centrifugation for 30 minutes at 33,000 g (Beckman JA 25.50).

Filtered supernatant was loaded onto a 5 ml HisTrap FF column (GE Healthcare) pre-equilibrated with lysis buffer. An Ӓkta Purifier FPLC was used to wash the column with 4% elution buffer (50 mM MES pH 6.5, 500 mM Imidazole, 300 mM NaCl) until absorbance at 280 nm reached a plateau. The wash was followed by a 50 mL gradient of lysis buffer to elution buffer at 1 mL/min. Protein elution continued by following the absorbance at 280 nm until all protein was eluted from the column, and the main peak collected for further purification. Fractions containing PIV eluted from the HisTrap column were pooled and directly loaded on a 5 mL Strep-Tactin XT gravity flow column. The column was pre-equilibrated with wash buffer (50 mM NaPO4, 150 mM NaCl, pH 5.8), then after sample was loaded the column was washed with 5 X 1 CV of wash buffer. The column was equilibrated with 1 CV of elution buffer (50 mM NaPO4, 150 mM NaCl, 50 mM Biotin, pH 6.3), and the protein eluted with 1.6 CV of elution buffer. The elution fraction containing pure PIV, confirmed by SDS PAGE, was concentrated to ∼10 mg mL^-1^ using a 30 kDa MWCO spin concentrator. Protein was flash frozen in liquid nitrogen and stored at -80°C for further experiments. Protein concentration was calculated from A280 readings using the extinction coefficient from Expasy protparam^38^ and a Denovix DS series instrument, or NanoDrop.

### PIV Activation

Aliquots of PIV were thawed and diluted with ∼5X with either wash buffer (50 mM NaPO4, 150 mM NaCl, pH 5.8) or assay buffer (50 mM Tris, 200 mM NaCl, pH 8.5) and incubated at room temperature for 4 days. Activation was followed by SDS-PAGE and further analysed by intact protein mass spectrometry.

### Cleavage of Cap sequence peptides containing lysine or ornithine

PIV was incubated with ornithine peptide for 1 or 24 hours (100 nM PIV, 100 uM APGTPLQVGVGLOTATPE, in assay buffer). One hundred microliter of reaction were collected from and quenched with an equal volume of 2% trifluoroacetic acid (TFA). The quenched sample was centrifuged at 16000 x g for 15 min at RT to remove precipitated protein. Caffeine, as an internal standard, was spiked into all samples to a final concentration of 10 μM before LC-MS analysis. Ten microliters of each sample were injected into XSelect PREMIER^™^ HSS T3 (2.5 μm, 4.6 x 50 mm) column. The gradient used 0.1% formic acid in water (A) and 0.1% formic acid in acetonitrile (B) as follows: 0-2 min: 1% B; 2-9 min: 1%-99% B in a linear gradient; 9-11 min: 99% B; 11-14 min: 1% B) at a flow rate of 0.4 mL/min. Elution of peptides of interested were carried out using the selected ion recording (SIR) function in the instrument (all m/z monitored in SIR channels are listed on Table S6). Analytes were detected using a ACQUITY QDa detector (Waters). SIR peaks were integrated, peak areas were normalized as a ratio to the peak area of caffeine to correct for discrepancies in injection volume. All experiments were performed with three biological replicates.

### PIV inhibition by Cap sequence peptides containing lysine or ornithine

PIV activity was measured using 10 nM PIV and T-GPK-NA at final concentrations of 0.010, 0.025, 0.050, 0.10, 0.20, 0.30, 0.40, or 0.50 mM. Assays contained either K-peptide (APGTPLQVGVGLKTATPE) or O-peptide (APGTPLQVGVGL(ornithine)TATPE) at a final concentration of 100, 150, or 200 µM.

### PIV Activity Assays

PIV activity was determined using N-(p-tosyl)-Gly-Pro-Lys-4-nitroanilide (T-GPK-NA; Sigma-Aldrich, T6140). Reactions were performed in in 50 mM Tris, 150 mM NaCl, pH 8.5 at 25°C. The release of p-nitroanilide was monitored by absorbance at 405 nM using a POLARstar Omega microplate reader (BMG LABTECH). Absorbance values were pathlength-corrected using the instrument’s built-in water-peak correction. Corrected absorbance values were converted to p-nitroaniline (pNA) concentrations using a pNA standard curve. The reaction was followed in 30-60 second intervals for 15-30 minutes and Initial reaction rates calculated from the linear portion of the product-formation curve.

### PIV Inhibition by PMSF and TLCK

PIV activity was measured using 10 nM PIV and T-GPK-NA at final concentrations of 0.010, 0.025, 0.050, 0.10, 0.20, 0.30, 0.40, or 0.50 mM. Reactions contained phenylmethylsulfonyl fluoride (PMSF) at 0 or 1 mM and Nα-tosyl-L-lysine chloromethyl ketone (TLCK) at 0, 10, or 50 µM.

### PIV-TLCK Jump Dilution

PIV (5 µM) was incubated with 10 µM TLCK for 0, 1, 2, 4, 8, or 16 h. The incubation mixtures were diluted to 2nM PIV and mixed in equal volume with 200 uM T-GPK-NA for a final assay concentration of 1 nM PIV, 100 µM T-GPK-NA, and 2 nM TLCK.

### PIV-dependent activation of PaAP

PaAP activity was determined using Leu-pNA (Bachem AG, 4001072.0005) as a substrate. Reactions were performed in in 50 mM Tris, 150 mM NaCl, pH 8.5 at 25°C. The release of p-nitroanilide was monitored by absorbance at 405 nM using a POLARstar Omega microplate reader (BMG LABTECH). Reactions contained 50 nM PaAP, 0, 0.5, 5, or 50 nM PIV and 1 mM Leu-pNA.

### Proteomics analysis of extracellular proteins and data analysis

30 mL of cell-free supernatant was collected from a 50 mL culture of P. aeruginosa PA14 strain grown in M9 minimum medium supplied with 0.5% casein for 24 hours and dried with freezer dryer, followed by resuspending in 1.5 mL of sterile PBS. 545 μL of 75% saturated trichloroacetic acid (TCA) was added to the supernatant to reach the final concentration of 20%, and the mixture was vortexed and incubated at -20℃. Precipitates were collected by centrifugating at 1600 g for 30 minutes, followed by washing with 0.5 mL of pre-chilled acetone for three times and drying under air flow.

Sample preparation with S-Trap columns was conducted according to the instructions from ProtiFi^39^. Briefly, dried samples were resuspended in lysis buffer (50 mM Tris-HCl, 10% SDS (w/v), pH 8.5), and protein concentrations were determined with BCA assay utilising BSA as standard and adjusted to 2.5 g/L. Proteins were reduced by adding TCEP to 5 mM and incubating at 55℃ for 15 minutes, followed by incubating with 20 mM IAA in the dark at 20℃ for 30 minutes in the dark. Samples were acidified using 2.5 μL of 37.5% phosphoric acid and 165 μL of binding/washing buffer (100 mM Tris-HCl in methanol (90%, v/v)) before loading to a S-Trap column. The column was washed with 150 μL of binding/washing buffer for another three times, and 10 μg of sequencing grade trypsin dissolved in 50 mM ammonium bicarbonate (AmBic) was added to the column for the overnight trypsin digestion at 37℃.

Peptides were eluted with three elution buffers in order: 40 μL of 50 mM Tris-HCl in water, 0.2% formic acid in water, and 50% acetonitrile in water. All elutes were pooled, dried with SpeedVac, and resuspended in loading buffer (0.05% TFA in water) to 1 μg/μL.

Samples were subjected to LC-MS/MS using an Ultimate 3000 RSLC (Thermo Scientific) coupled to an Orbitrap Fusion Lumos mass spectrometer (Thermo Scientific) in data-dependent acquisition (DDA) mode. Raw data was imported in MaxQuant (v 2.6.7.0). All settings were set to default unless noted. Carbamidomethylation (C) was set as a fixed modification, and oxidation on methionine was set as a variable modification, which were also used for protein quantification. Trypsin was selected as digestion enzyme, and the maximum number of missing cleavage site was set to 1. Data were searched against *Pseudomonas aeruginosa* UCBPP-PA14 protein database from Uniprot (acquired on May 17^th^, 2024), and the false discovery rate (FDR) of protein group identification was set to 0.01 based on the search in the reverse decoy database generated by MaxQuant^40^. Label-free quantification (LFQ) algorithm was used for relative quantification.

### Crystallisation, data collection and structure determination

Crystals grew readily in several commercial screens provided by Molecular Dimensions. Crystals were grown at 18°C using sitting drop vapour diffusion technique, with a drop size of 0.3 μl in a ratio of 1:1, reservoir:protein solution, using ATI gryphon. PIV_S409A_ was used at a concentration of 12.3 mg/ml for all crystallisation experiments. Initially, crystals diffracted to lower resolution, which was improved upon using random matrix microseeding^41^ (rMMS) on the Douglas Instruments Oryx4. rMMS trials were performed using a 3:2:1 ratio of protein:reservoir:seed stock, were initial seed stock (Screen: BCS, Condition: G9) was diluted 1 in 10 prior to use. Seeding resulted in improved crystal growth and diffraction quality across several conditions. Larger dagger-shaped crystals of PIV_S409A_, grown in the BCS D1 following seeding, were used to collect the high-resolution data (2.17 Å). A full list of crystallisation conditions can be found in Supplementary table 4. Crystals were transferred tp mother liquor supplemented with 30 % Ethylene glycol prior to being looped and flash cooled in liquid nitrogen, for storage until data collection.

Diffraction data were collected at the Diamond Light Source (Oxford, UK) on beamline I24. Single wavelength diffraction experiments was performed (Wavelength: 0.97 Å, Detector resolution: 2.5 Å, Exposure: 0.005 s, Transmission: 20%, Beamsize: 50 x 50 µm. Two datasets (3600 images at 0.1° oscillation) were collected at different sites using a single crystal. The automated pipeline mulit.xia2 was used for data reduction and processing, combining datasets for improved crystallographic statistics. The structure of PIV_S409A_ was solved by molecular replacement, using Phenix.phaser, with 4 copies of the AlphaFold model (acquired from Uniprot AF-Q02SZ7-F1). Subsequent model building and refinement were performed iteratively in Coot and Phenix. Crystallographic and refinement data are shown in Table S5.

### NanoDSF and dynamic light-scattering analysis

Mature WT PIV and PIV_S409A_ were diluted to 1 mg/ml in DSF buffer (50 mM NaPO4 pH 6.5, 150 mM NaCl) and loaded into standard capillaries. Nano-differential scanning fluorimetry (nanoDSF) and dynamic light scattering (DLS) measurements were performed using a Prometheus Panta instrument (NanoTemper Technologies). For nanoDSF analysis, intrinsic protein fluorescence at 330 and 350 nm was recorded during heating from 25 to 99 at a rate of 1 °C min⁻¹. Thermal unfolding was monitored using the A350/A330 fluorescence ratio, and apparent melting temperatures were determined from the corresponding first-derivative maxima. DLS measurements were performed at 25 °C using 10 acquisitions. Intensity-weighted particle-size distributions, hydrodynamic radii and polydispersity values were calculated using the manufacturer’s analysis software. Measurements were performed in triplicate, and results are reported as the mean ± SD.

### Intact protein mass analysis by LC-MS

The protein sample (20μL, 1 μM) was desalted on-line through a MassPrep On-Line Desalting Cartridge 2.1 Å∼ 10mm, using a Waters Acquity H-class HPLC, eluting at 200 μL/min, with an increasing acetonitrile concentration (2% acetonitrile, 98% aqueous 1% formic acid to 98% acetonitrile, 2% aqueous 1% formic acid) and delivered to a Waters Xevo G2XS electrospray ionisation mass spectrometer operated with positive polarity in sensitivity mode. Intermittently a lockspray signal using Leucine Enkephalin was measured and a mass correction was applied. An envelope of multiply charged signals was acquired between m/z 500–2500 and deconvoluted using MaxEnt1 software to give the molecular mass of the protein. For experiments in the presence of TLCK, 27 μM PIV was incubated overnight with 100 μM TLCK and submitted to analysis using the method described above.

### Gel Analysis and Data fitting

SDS-PAGE gel images were analysed in ImageJ^42^ (FIJI software) using the programs gel analysis tool. Gel bands were assigned manually with the rectangle selection tool encompassing different protein species. For band quantification, total protein in the lane was set to 100%, and each band intensity is reported as a fraction of this. Intensity values (arbitrary units, AU) were exported, normalised for total protein in the lane, plotted and analysed using GraphPad Prism.

## Statistical analysis

All data are presented as mean ± standard error of the mean and were obtained from ≥3 independent experiments with total sample numbers provided in the figure legends. Statistical significance was evaluated with GraphPad Prism software, using a one-sample t-test. Significant difference assessed through an unpaired t-test, ****P < 0.0001.

## Data availability

Structural data were deposited in the PDB and are available under accession numbers 31RM. All other data are contained in the main manuscript and supplementary information. Raw data accompanying figures is available for download.^30^ All materials and reagents are available from the corresponding authors. The mass spectrometry proteomics data have been deposited to the ProteomeXchange Consortium via the PRIDE partner repository with the dataset identifier PXD083431 and 10.6019/PXD083431.

## Acknowledgements

We thank Sally Shirran from the Mass Spectrometry Facility in St Andrews for performing secreted proteomics and intact protein mass experiments.

C.J.H., C.H. were funded by the Wellcome Trust (210486/Z/18/Z), C.M.C. and C.J.H. were funded by the Biotechnology and Biological Sciences Research Council (BB/Y005333/1). Q.G. was funded by Tenovus Research Scotland (Grant: T22-739).

## Conflic of interest

The authors declare that they have no conflicts of interest with the contents of this article.

## Author contributions

### Conceptualisation and study design

C.J.H., and C.M.C.

### Experiments

C.J.H. performed the crystallization and structural work and contributed to biochemical experiments including protein production (PIV S409A) and initial activity assays. C.H optimised recombinant production of WT PIV and performed activity assays as well as characterisation of inhibitors. Q.G. carried out proteomics experiments of secreted proteins.

### Methodology and supervision

C.M.C supervised the overall project.

### Data analysis and interpretation

C.J.H., C.H., Q.G., and C.M.C. were involved in analysing and interpreting data.

### Writing

C.J.H., and C.M.C. wrote the initial draft, and all authors contributed equally to the final manuscript draft, reviewing, editing and approving the final manuscript.

## Notes

### Competing Interest Statement

The authors have declared no competing interest.

## References

1 Zhao, T. H. et al. Extracellular aminopeptidase modulates biofilm development of Pseudomonas aeruginosa by affecting matrix exopolysaccharide and bacterial cell death. Environmental Microbiology Reports 10, 583–593, doi:10.1111/1758-2229.12682 (2018).

2 Galdino, A. C. M., Branquinha, M. H., Santos, A. L. S. & Viganor, L. in *Pathophysiological Aspects of Proteases* Ch. Chapter 16, 381–397 (2017).

3 Hennemann, L. C. & Nguyen, D. LasR-regulated proteases in acute vs. chronic lung infection: a double-edged sword. Microb Cell 8, 161–163, doi:10.15698/mic2021.07.755 (2021).

4 Cigana, C. et al. Pseudomonas aeruginosa Elastase Contributes to the Establishment of Chronic Lung Colonization and Modulates the Immune Response in a Murine Model. Frontiers in Microbiology Volume 11 - 2020, doi:10.3389/fmicb.2020.620819 (2021).

5 Strateva, T. & Mitov, I. Contribution of an arsenal of virulence factors to pathogenesis of Pseudomonas aeruginosa infections. Annals of Microbiology 61, 717–732, doi:10.1007/s13213-011-0273-y (2011).

6 Bleves, S. et al. Protein secretion systems in Pseudomonas aeruginosa: A wealth of pathogenic weapons. International Journal of Medical Microbiology 300, 534–543, doi:10.1016/j.ijmm.2010.08.005 (2010).

7 Kipnis, E., Sawa, T. & Wiener-Kronish, J. Targeting mechanisms of Pseudomonas aeruginosa pathogenesis. Medecine Et Maladies Infectieuses 36, 78–91, doi:10.1016/j.medmal.2005.10.007 (2006).

8 Saint-Criq, V. et al. Pseudomonas aeruginosa LasB protease impairs innate immunity in mice and humans by targeting a lung epithelial cystic fibrosis transmembrane regulator–IL-6–antimicrobial–repair pathway. Thorax 73, 49, doi:10.1136/thoraxjnl-2017-210298 (2018).

9 Ołdak, E. & Trafny Elżbieta, A. Secretion of Proteases by Pseudomonas aeruginosa Biofilms Exposed to Ciprofloxacin. Antimicrobial Agents and Chemotherapy 49, 3281–3288, doi:10.1128/aac.49.8.3281-3288.2005 (2005).

10 Axelrad, I. et al. Extracellular proteolytic activation of Pseudomonas aeruginosa aminopeptidase (PaAP) and insight into the role of its non-catalytic N-terminal domain. PLoS One 16, e0252970, doi:10.1371/journal.pone.0252970 (2021).

11. Engel, L. S., Hill, J. M., Caballero, A. R., Green, L. C. & O’Callaghan, R. J. Protease IV, a Unique Extracellular Protease and Virulence Factor from *Pseudomonas aeruginosa* *. Journal of Biological Chemistry 273, 16792–16797, doi:10.1074/jbc.273.27.16792 (1998).

12 Giansanti, P., Tsiatsiani, L., Low, T. Y. & Heck, A. J. R. Six alternative proteases for mass spectrometry–based proteomics beyond trypsin. Nature Protocols 11, 993–1006, doi:10.1038/nprot.2016.057 (2016).

13 Caballero, A., Thibodeaux, B., Marquart, M., Traidej, M. & O’Callaghan, R. Pseudomonas Keratitis: Protease IV Gene Conservation, Distribution, and Production Relative to Virulence and Other Pseudomonas Proteases. Investigative Ophthalmology & Visual Science 45, 522–530, doi:10.1167/iovs.03-1050 (2004).

14 Engel, L. S. et al. Pseudomonas aeruginosa protease IV produces corneal damage and contributes to bacterial virulence. Investigative Ophthalmology & Visual Science 39, 662–665 (1998).

15 Engel, L. S. et al. Pseudomonas deficient in protease IV has significantly reduced corneal virulence. Investigative Ophthalmology & Visual Science 38, 1535–1542 (1997).

16 O’Callaghan, R., Caballero, A., Tang, A. & Bierdeman, M. Pseudomonas aeruginosa Keratitis: Protease IV and PASP as Corneal Virulence Mediators. Microorganisms 7, doi:10.3390/microorganisms7090281 (2019).

17 Oh, J., Li, X.-H., Kim, S.-K. & Lee, J.-H. Post-secretional activation of Protease IV by quorum sensing in Pseudomonas aeruginosa. Scientific Reports 7, 4416, doi:10.1038/s41598-017-03733-6 (2017).

18 Li, X. H. & Lee, J. H. Quorum sensing-dependent post-secretional activation of extracellular proteases in Pseudomonas aeruginosa. J Biol Chem 294, 19635–19644, doi:10.1074/jbc.RA119.011047 (2019).

19. Daboor, S. M. et al. Inhibition of *Pseudomonas aeruginosa-*secreted protease IV reduces lung inflammation. bioRxiv, 2026.2007.2027.739189, doi:10.64898/2026.07.27.739189 (2026).

20. Traidej, M., Marquart, M. E., Caballero, A. R., Thibodeaux, B. A. & O’Callaghan, R. J. Identification of the Active Site Residues of *Pseudomonas aeruginosa* Protease IV: IMPORTANCE OF ENZYME ACTIVITY IN AUTOPROCESSING AND ACTIVATION *. Journal of Biological Chemistry 278, 2549-2553, doi:10.1074/jbc.M208973200 (2003).

21 Jumper, J. et al. Highly accurate protein structure prediction with AlphaFold. Nature 596, 583–589, doi:10.1038/s41586-021-03819-2 (2021).

22 Abramson, J. et al. Accurate structure prediction of biomolecular interactions with AlphaFold 3. Nature 630, 493–500, doi:10.1038/s41586-024-07487-w (2024).

23 Krissinel, E. & Henrick, K. Inference of macromolecular assemblies from crystalline state. J Mol Biol 372, 774–797, doi:10.1016/j.jmb.2007.05.022 (2007).

24 Holm, L. & Sander, C. DALI - A NETWORK TOOL FOR PROTEIN-STRUCTURE COMPARISON. Trends in Biochemical Sciences 20, 478–480, doi:10.1016/s0968-0004(00)89105-7 (1995).

25 van Kempen, M. et al. Fast and accurate protein structure search with Foldseek. Nat. Biotechnol. 42, 243–246, doi:10.1038/s41587-023-01773-0 (2024).

26 van der Hoeven, L. R., Lechner, M., Hernandez-Rollan, C., Batth, T. S. & Olsen, J. V. Comparative Analysis of Lysine-Specific Peptidases for Optimizing Proteomics Workflows. Journal of Proteome Research 25, 1176–1183, doi:10.1021/acs.jproteome.5c00872 (2025).

27 Khavrutskii, I. V., Compton, J. R., Jurkouich, K. M. & Legler, P. M. Paired Carboxylic Acids in Enzymes and Their Role in Selective Substrate Binding, Catalysis, and Unusually Shifted pK(a) Values. Biochemistry 58, 5351–5365, doi:10.1021/acs.biochem.9b00429 (2019).

28 Parker, D. S. et al. Retrospective analysis of secreted PrpL protease activity in clinical isolates of Pseudomonas aeruginosa and its association with corneal tissue damage. Frontiers in Microbiology Volume 17 - 2026, doi:10.3389/fmicb.2026.1824817 (2026).

29 Harding, C. J., Bischoff, M., Bergkessel, M. & Czekster, C. M. An anti-biofilm cyclic peptide targets a secreted aminopeptidase from P. aeruginosa. Nature Chemical Biology 19, 1158–1166, doi:10.1038/s41589-023-01373-8 (2023).

30. Gong, Q., PRIDE, Extracellular proteome (P. aeruginosa PA14 strain), Project accession: PXD083431. DOI: 10.6084/m9.figshare.26380666.v2

31 Ohnishi, Y. et al. Neutron and X-ray crystallographic analysis of Achromobacter protease I at pD 8.0: Protonation states and hydration structure in the free-form. Biochimica et Biophysica Acta (BBA) - Proteins and Proteomics 1834, 1642–1647, 10.1016/j.bbapap.2013.05.012 (2013).

32 Tsunasawa, S., Masaki, T., Hirose, M., Soejima, M. & Sakiyama, F. The primary structure and structural characteristics of Achromobacter lyticus protease I, a lysine-specific serine protease. J Biol Chem 264, 3832–3839 (1989).

33 Asztalos, P., Müller, A., Hölke, W., Sobek, H. & Rudolph, M. G. Atomic resolution structure of a lysine-specific endoproteinase from Lysobacter enzymogenes suggests a hydroxyl group bound to the oxyanion hole. Acta Crystallogr D Biol Crystallogr 70, 1832–1843, doi:10.1107/s1399004714008463 (2014).

34 Fersht, A. R. & Renard, M. pH Dependence of chymotrypsin catalysis. Appendix. Substrate binding to dimeric α-chymotrypsin studied by x-ray diffraction and the equilibrium method. Biochemistry 13, 1416–1426, doi:10.1021/bi00704a016 (2002).

35 Malthouse, J. P. G. Kinetic Studies of the Effect of pH on the Trypsin-Catalyzed Hydrolysis of N-α-benzyloxycarbonyl-l-lysine-p-nitroanilide: Mechanism of Trypsin Catalysis. ACS Omega 5, 4915–4923, doi:10.1021/acsomega.9b03750 (2020).

36 Plotnick, M. I. et al. Heterogeneity in Serpin−Protease Complexes As Demonstrated by Differences in the Mechanism of Complex Breakdown. Biochemistry 41, 334–342, doi:10.1021/bi015650+ (2001).

37 Zhu, X., Ohta, Y., Jordan, F. & Inouye, M. Pro-sequence of subtilisin can guide the refolding of denatured subtilisin in an intermolecular process. Nature 339, 483–484, doi:10.1038/339483a0 (1989).

38. Gasteiger, E. H., C. Gattiker, A. Duvaud, S. Wilkins, M. R. Appel, R. D. Bairoch, A. in The Proteomics Protocols Handbook (ed John M. Walker) 571-607 (Humana Press, 2005).

39 HaileMariam, M. et al. S-Trap, an Ultrafast Sample-Preparation Approach for Shotgun Proteomics. J Proteome Res 17, 2917–2924, doi:10.1021/acs.jproteome.8b00505 (2018).

40 Tyanova, S., Temu, T. & Cox, J. The MaxQuant computational platform for mass spectrometry-based shotgun proteomics. Nature Protocols 11, 2301–2319, doi:10.1038/nprot.2016.136 (2016).

41 Stewart, P. D. S., Kolek, S. A., Briggs, R. A., Chayen, N. E. & Baldock, P. F. M. Random Microseeding: A Theoretical and Practical Exploration of Seed Stability and Seeding Techniques for Successful Protein Crystallization. Crystal Growth & Design 11, 3432–3441, doi:10.1021/cg2001442 (2011).

42 Schneider, C. A., Rasband, W. S. & Eliceiri, K. W. NIH Image to ImageJ: 25 years of image analysis. Nature Methods 9, 671–675, doi:10.1038/nmeth.2089 (2012).

