## Supplementary material for "Structural basis for autoinhibition and self-activation in the *Pseudomonas aeruginosa* virulence factor protease IV (PIV)": SI

### Supplementary Information Figures

**Figure S1: Biophysical characterisation of mature WT PIV and PIV S409A by nano-differential scanning fluorimetry (nanoDSF) and dynamic light scattering (DLS).**

Measurements were acquired using a NanoTemper Prometheus Panta. Tabulated values are presented as the mean ± SD of triplicate measurements and are reported to two decimal places; plots show one representative measurement. A, C, Thermal unfolding profiles of mature wild-type PIV (A) and PIV S409A (C), showing the intrinsic fluorescence ratio (F350/F330) as a function of temperature and its corresponding first derivative. B, D, Intensity-weighted DLS particle-size distributions for mature wild-type PIV (B) and PIV S409A (D). Wild-type PIV was dominated by a single, monodisperse population with a hydrodynamic radius consistent with monomeric protein. PIV S409A contained a similarly sized monodisperse population (peak 1) and a second, broader population of larger particles (peak 2). Because light-scattering intensity increases strongly with particle size, larger particles contribute disproportionately to an intensity-weighted DLS distribution. The relative peak intensities should therefore not be interpreted directly as the relative numbers or concentrations of particles in each population.

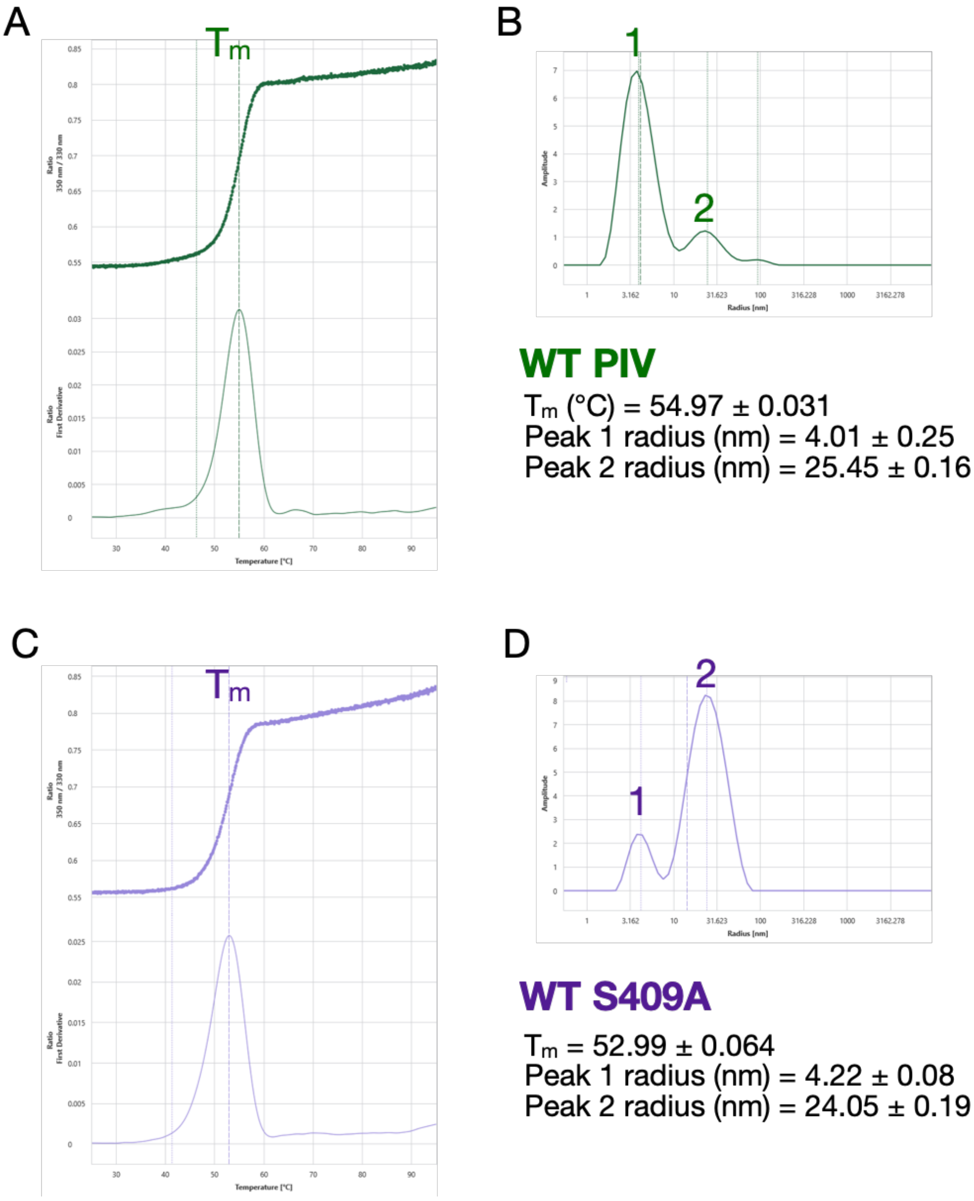

**Figure S2: SDS-PAGE depicting the purification of active site mutants of PIV.**

SDS-PAGE following purification of PIV active site mutants. The yellow arrow indicates the full-length PIV species at ~48 kDa. In this gel T = total protein, S = soluble protein loaded onto histrap column and B1&2 = protein eluted from the column using high imidazole elution buffer. $PIV_{S409A}$ was used for x-ray crystallography (indicated by yellow arrow).

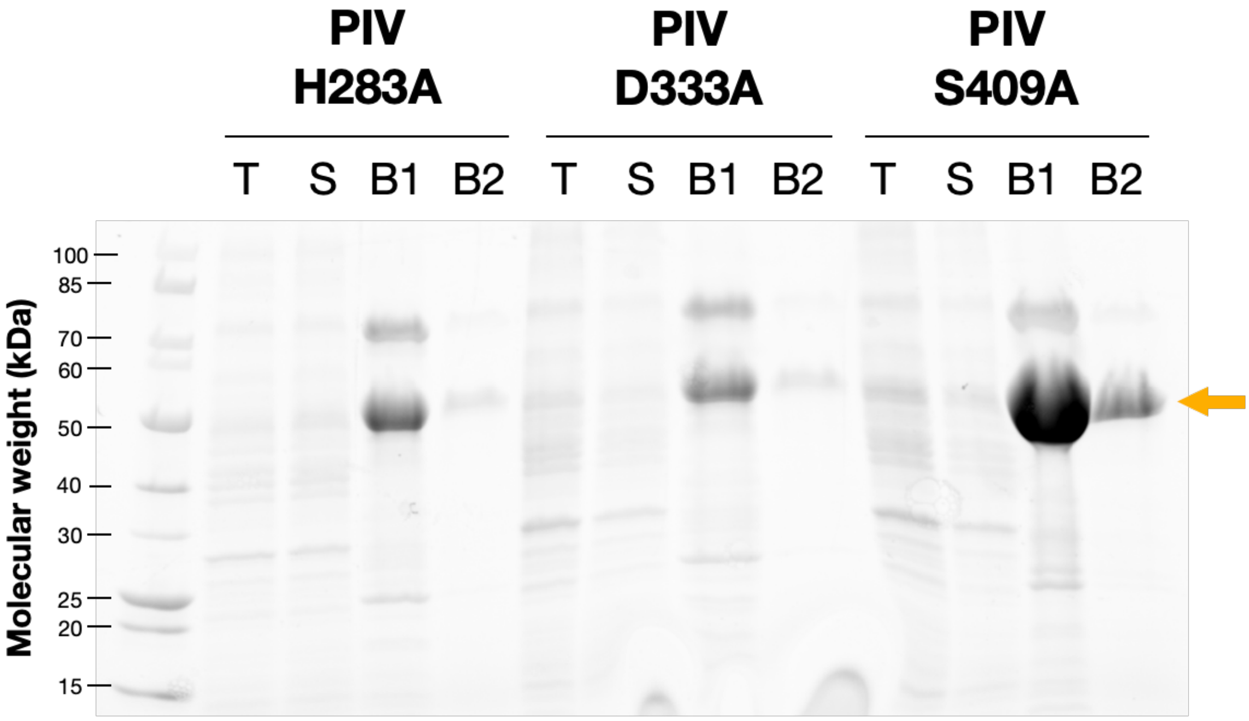

**Figure S3: Wild type PIV (WT) can be seen as a "3-protein intermediate" which undergoes self-cleavage to complete degradation during the purification.**

SDS-PAGE following purification of PIV variants at pH 8.5. WT PIV shows a pattern of self-cleavage in initial purification attempts - gel lane labelled as "WT B". The same degradation pattern is absent in the inactive variants H283A and D333A during identical purification steps. In this gel T = total protein, S = soluble protein loaded onto histrap column and B = protein eluted from the column using imidazole. The yellow arrow indicates the full-length PIV species.

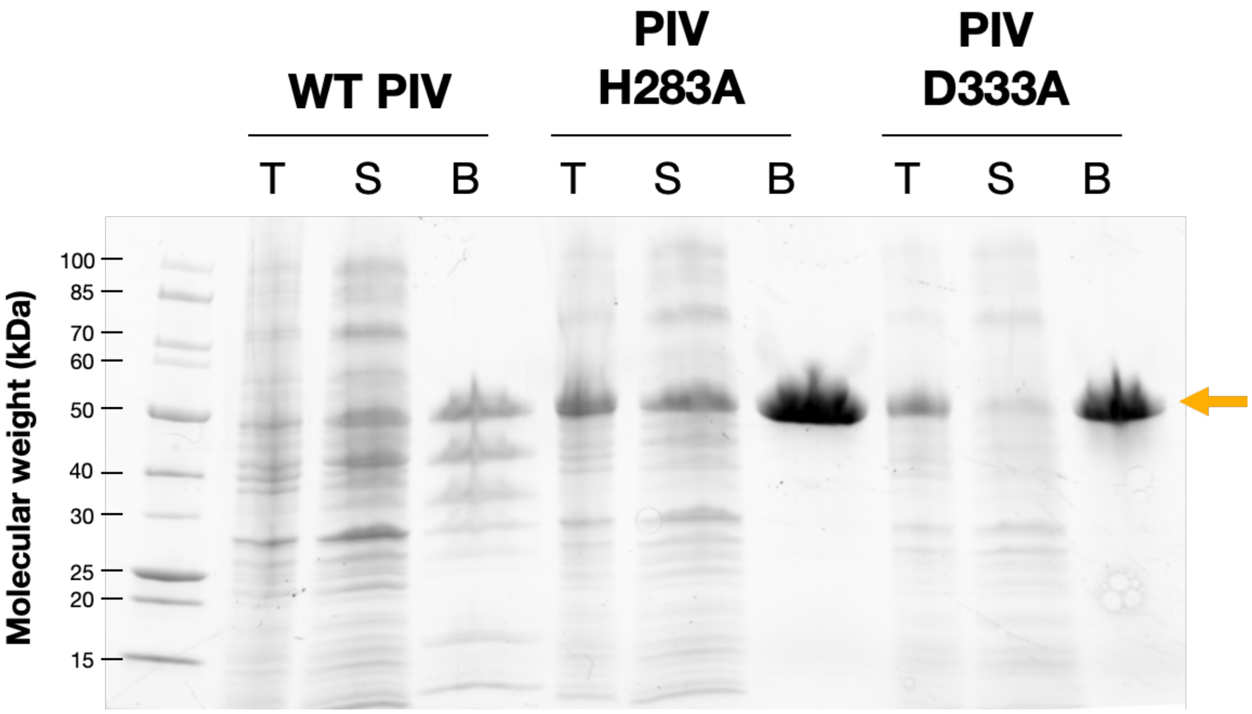

**Figure S4: Mapping of PIV lysine residues and list of cleavage sites**

Structure of PIV in ribbon form with lysine mapped. Lysine residues are coloured according to their location within the structure and their accessibility: magenta lys residues belong to the Cap-CUB assembly; green lys residues belong to the protease domain; and blue lys residues are inaccessible protease domain located residues. Below the structure is the sequence register +/- eight residues around the selective lysine residue cleavage site, colour matched to the structure.

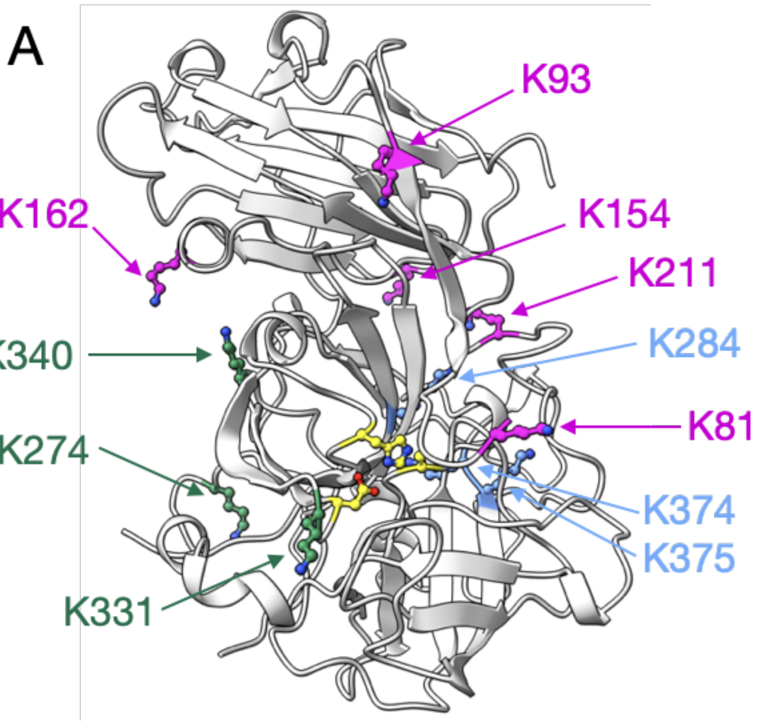

| | |
|---|---|
| K81 | ARAAPLAPKPGTPLQVG |
| K93 | PLQVGVGLKTATPEIDL |
| K154 | LLHFAGAGKEIFEASGK |
| K162 | KEIFEASGKDLSVNRPY |
| K211 | SYFADSLYKAGYRDGFG |
| K274 | DNATAAVAKMVFTSSAD |
| K284 | LLNNGNSPKRQLFWSAA |
| K331 | ANILHRDAKRDTLLLEL |
| K340 | RDTLLLELKRTPPAGVF |
| K374 | IHHPRGDAKKYSQGNVS |
| K375 | HHPRGDAKKYSQGNVSA |

| Cleavage sites | No-evidence for cleavage |
|---|---|
| GWSHPQFEKGAAPGASE – Streptag | PLQVGVGLKTATPEIDL |
| ARAAPLAPKPGTPLQVG - proposed | LLHFAGAGKEIFEASGK |
| KEIFEASGKDLSVNRPY | DNATAAVAKMVFTSSAD |
| SYFADSLYKAGYRDGFG | LLNNGNSPKRQLFWSAA |
| | ANILHRDAKRDTLLLEL |
| | RDTLLLELKRTPPAGVF |
| | IHHPRGDAKKYSQGNVS |
| | HHPRGDAKKYSQGNVSA |

#### Figure S5. Intact protein mass spectrometry of PIV variants and cleavage intermediates

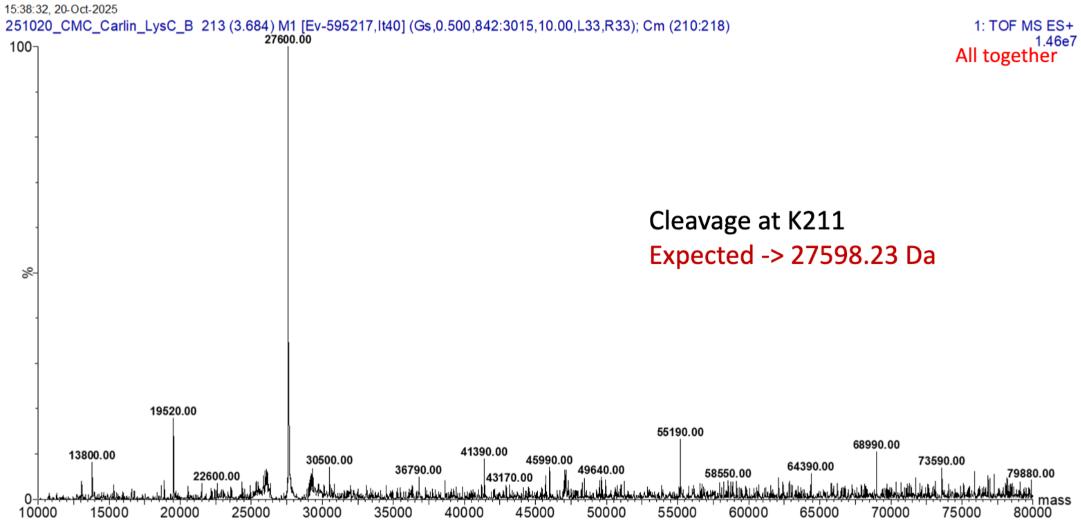

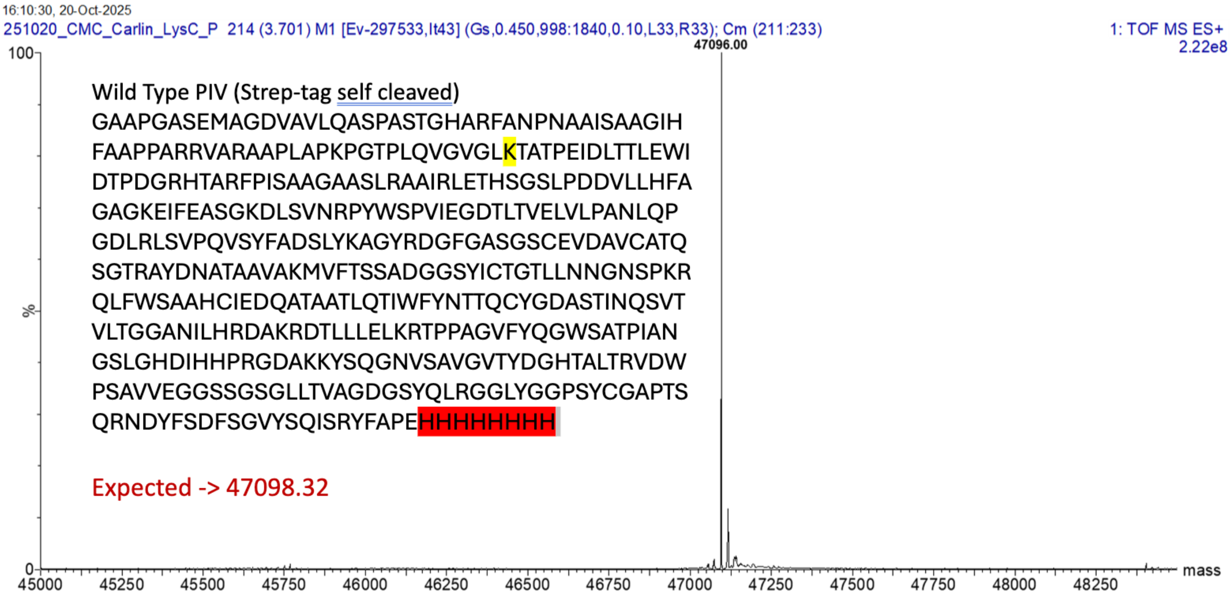

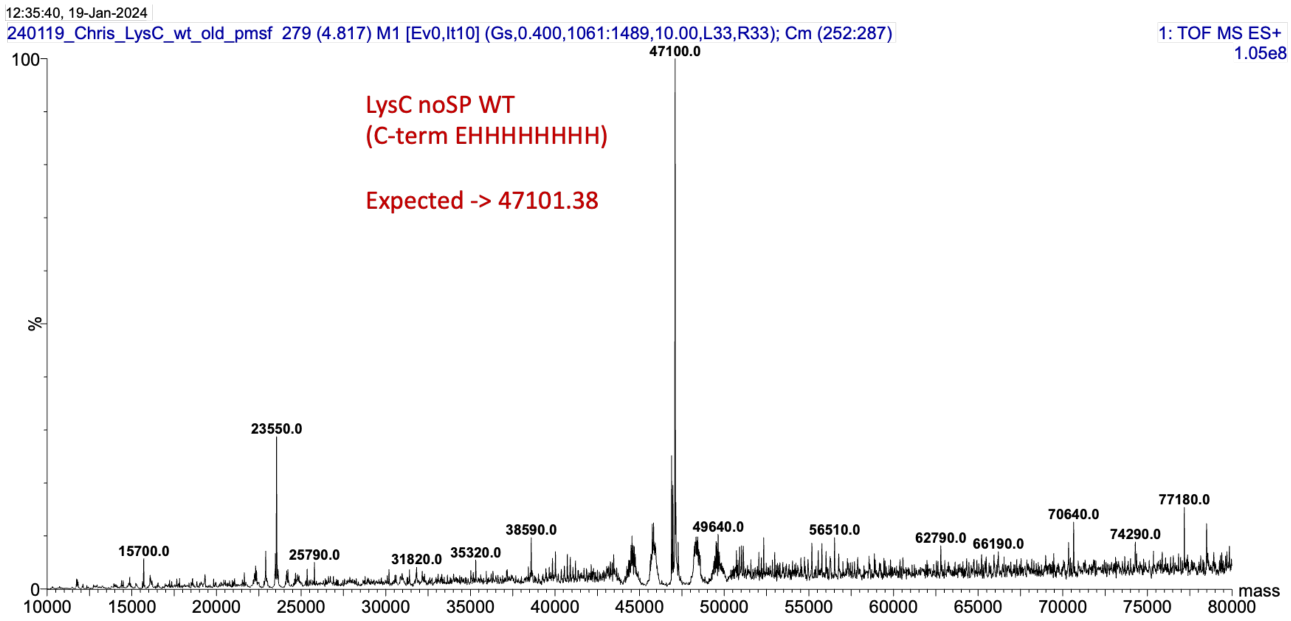

PIV purified in the presence of PMSF shows no covalent modification.

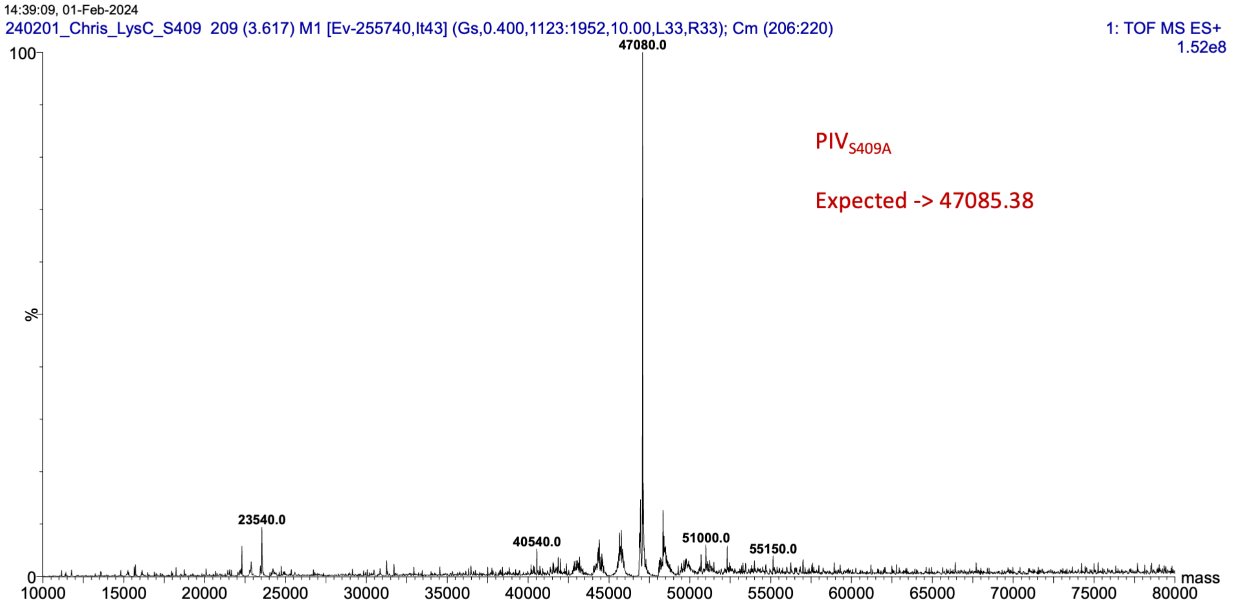

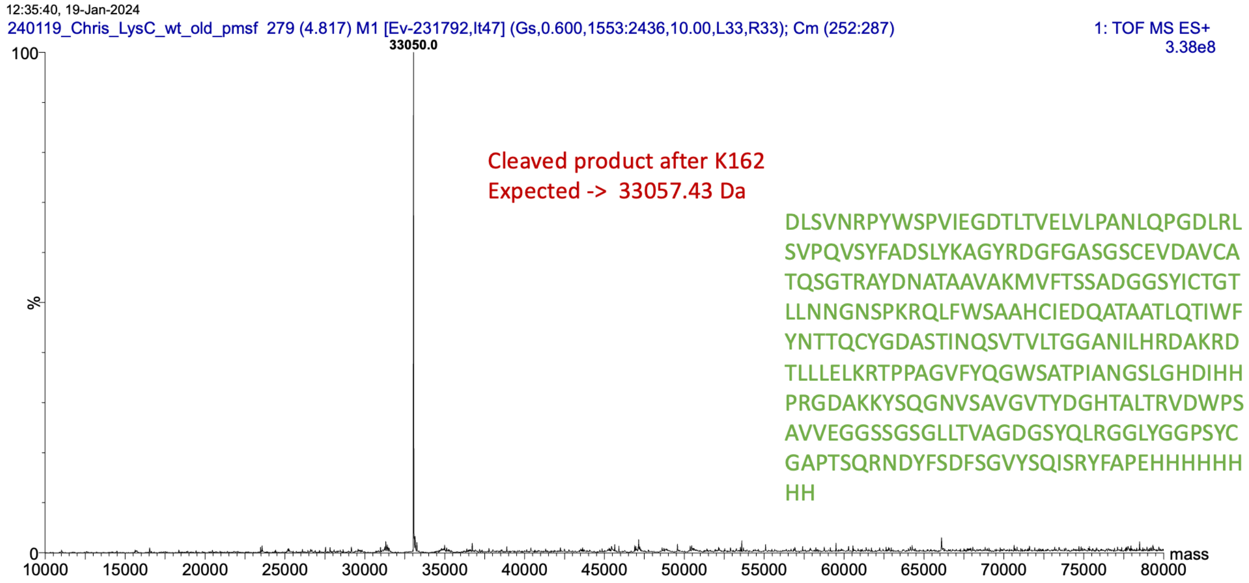

**Figure S6: PIV is covalently inhibited by TLCK**

a) Pre-incubation of PIV:TLCK at different ratios for 4h followed by jump dilution shows no activity recovery, indicative or an irreversible inhibition process. b) Intact protein mass spectrometry showing the formation of an additional adduct with a mass addition of 297 Da after overnight incubation with TLCK (27 µM PIV incubated with 100 µM TLCK), demonstrating incomplete covalent modification by TLCK, as the unmodified protein can still be seen (27592 Da).

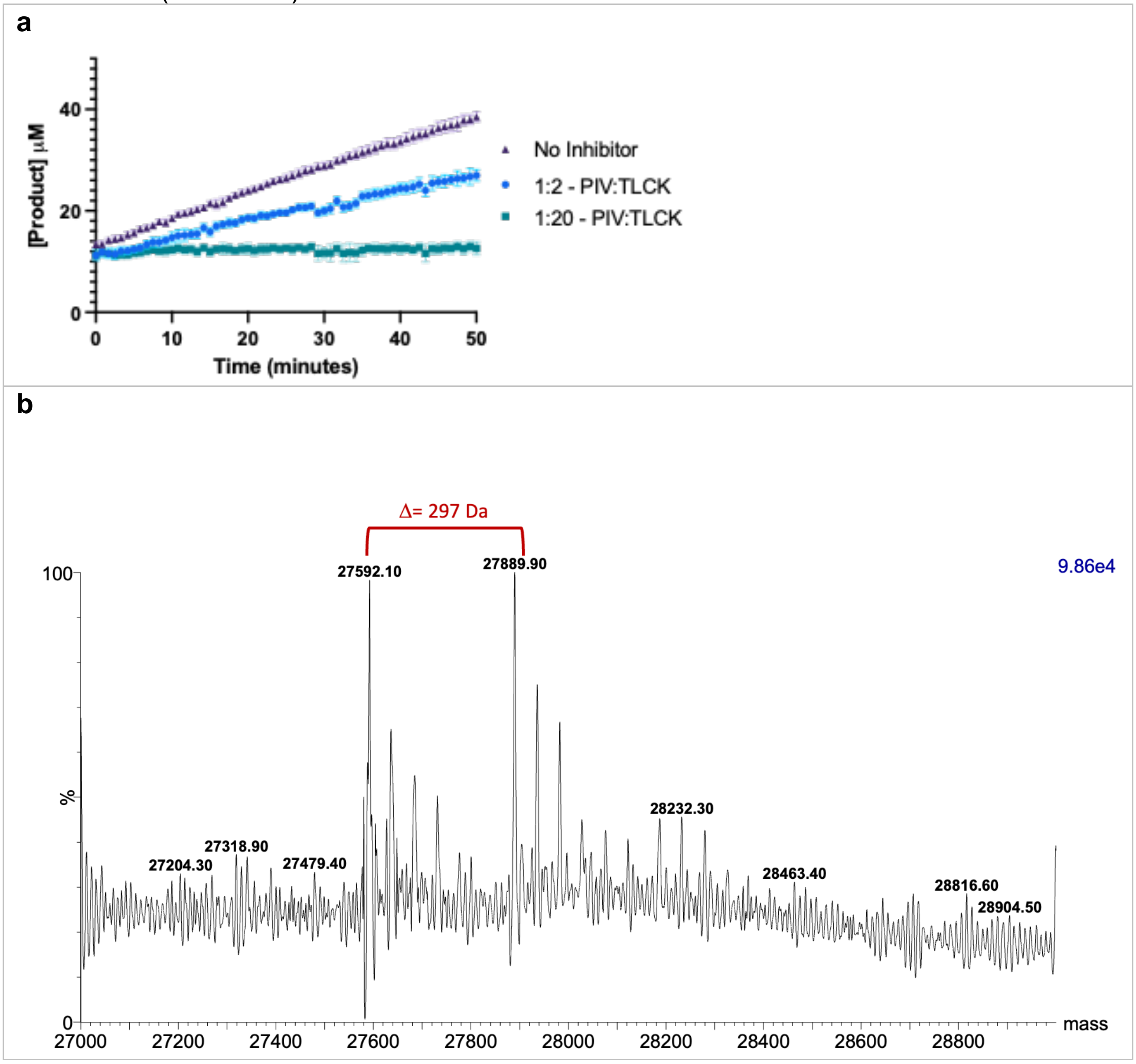

**Figure S7: Non-linearized data for Figure 4E and 4F.**

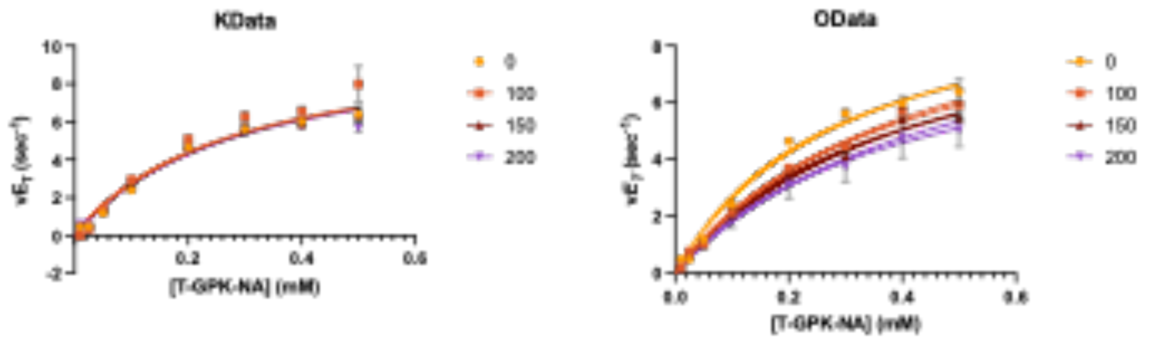

**Figure S8: Influence of peptides that contain the same sequence as PIV's cap in the reaction velocity.**

**A)** In the presence of excess PIV substrate, a modest effect can be seen in longer Cap sequences that contained lysine of ornithine. **B)** Lysine containing peptides are readily cleaved following incubation with PIV. Normalised peak areas (relative to caffeine as a standard) of the intact K- and O-peptides are shown after incubation in the presence (+ PIV) or absence (-PIV) of PIV. Bars represent the mean ± SD (n=3) of separate and independent reactions. **C)** Representative SIR (selected ion recording) chromatograms of the K- and O-peptides following incubation with or without PIV. Intact peptides, N-terminal products, and C-terminal products are shown in blue, green, and brown, respectively. SIR targets are shown in Table S6.

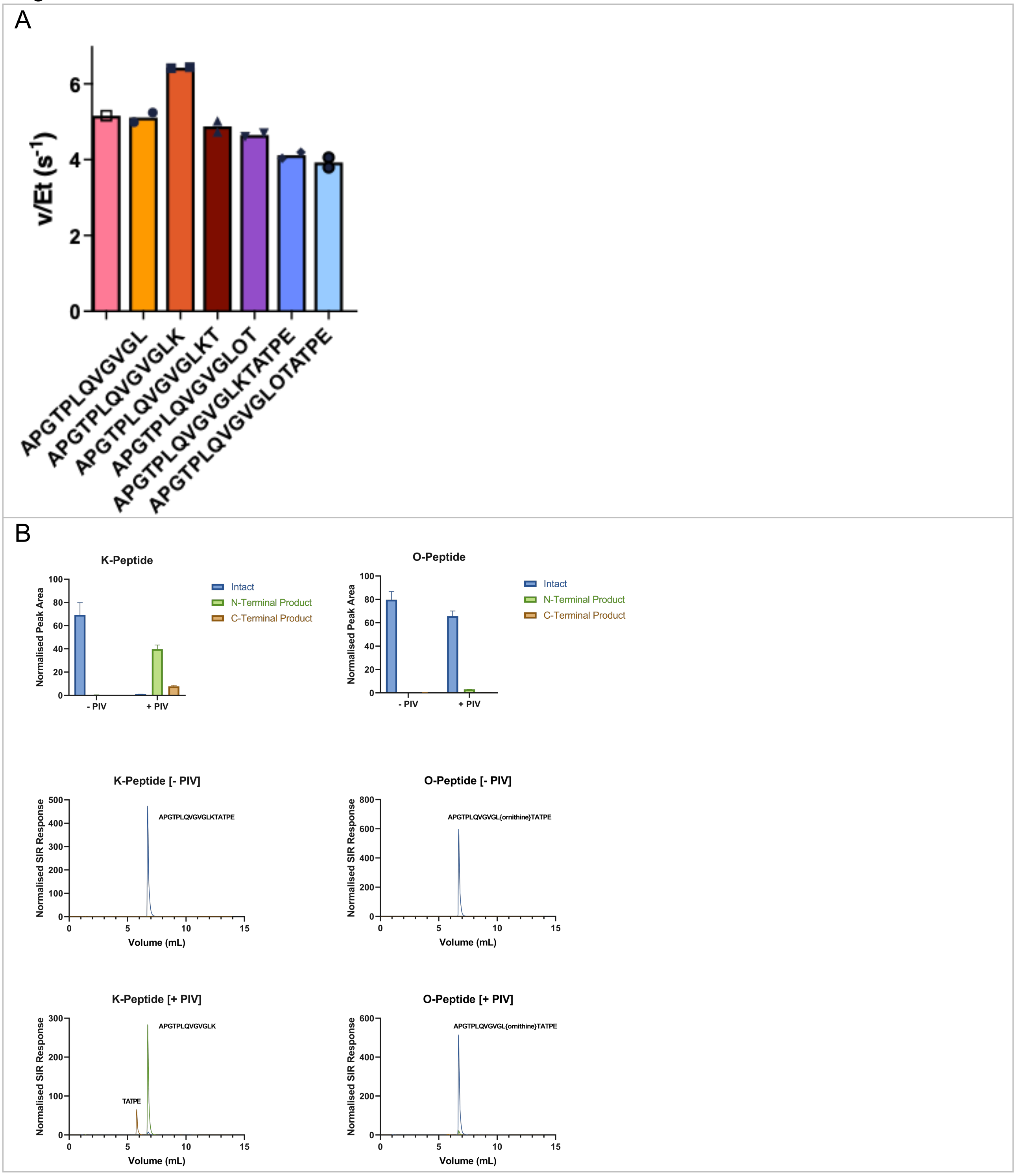

**Figure S9: Proteomic analysis of secreted proteins in P. aeruginosa PA14 cultures shows mature PIV and active PaAP are readily observed.**

Sequences in bold were identified in all three biological replicates, and residues with shading in PA14_09900 are identified by at least two replicates.

PaAP – last 24 residues were not observed in the enzyme form present in the supernatant

| | | | | | |
|---|---|---|---|---|---|
| 1 | MSNKNNLRYA | LGALALSVSA | ASLAAPSEAQ | QFTEFWTPGK | PNPSICK**SPL** |
| 51 | **LVSTPLGLPR** | **CLQASNVVKR** | **LQKLEDIASL** | **NDGNRAAATP** | **GYQASVDYVK** |
| 101 | **QTLQKAGYKV** | **SVQPFPFTAY** | **YPKGPGSLSA** | **TVPQPVTYEW** | **EKDFTYLSQT** |
| 151 | **EAGDVTAKVV** | **PVDLSLGAGN** | **TSTSGCEAED** | **FANFPAGSIA** | **LIQRGTCNFE** |
| 201 | **QKAENAAAAG** | **AAGVIIFNQG** | **NTDDRKGLEN** | **VTVGESYEGG** | **IPVIFATYDN** |
| 251 | **GVAWSQTPDL** | **QLHLVVDVVR** | K**KTETYNVVA** | **ETRRGNPNNV** | **VMVGAHLDSV** |
| 301 | **FEGPGINDNG** | **SGSAAQLEMA** | **VLLAKALPVN** | **KVRFAWWGAE** | **EAGLVGSTHY** |
| 351 | **VQNLAPEEKK** | KIK**AYLNFDM** | **IGSPNFGNFI** | **YDGDGSDFGL** | **QGPPGSAAIE** |
| 401 | **RLFEAYFRLR** | **GQQSEGTEID** | **FRSDYAEFFN** | **SGIAFGGLFT** | **GAEGLKTEEQ** |
| 451 | **AQKYGGTAGK** | **AYDECYHSKC** | **DGIANINQDA** | **LEIHSDAMAF** | **VTSWLSLSTK** |
| 501 | **VVDDEIAAAG** | **QK**AQSRSLQM | QKSASQIERW | GHDFIK | |

PIV – Cap and CUB domains not observed in the mature secreted protein, only peptides C-terminus from K211

| | | | | | |
|---|---|---|---|---|---|
| 1 | MHKRTYLNAC | LVLALAAGAS | QASAAPGASE | MAGDVAVLQA | SPASTGHARF |
| 51 | ANPNAATSAA | GIHFAAPPAR | RVARAAPLAP | KPGTPLQVGV | GLKTATPEID |
| 101 | LATLEWIDTP | DGRHTARFPI | SAAGAASLRA | AIRLETRSGS | LPDDVLLHFA |
| 151 | GAGKEIFEAS | GKDLSLNRPY | WSPVIEGDTL | TVELVLPANL | QPGDLRLSVP |
| 201 | QVSYFADSLY | K**AGYRDGFGA** | **SGSCEVDAVC** | **ATQSGTRAYD** | **NATAAVAKMV** |
| 251 | **FTSSADGGSY** | **ICTGTLLNNG** | **NSPK**RQLFWS | AAHCIEDQAT | AATLQTIWFY |
| 301 | NTTQCYGDAS | TINQSVTVLT | GGANILHRDA | K**RDTLLLELK** | R**TPPAGVFYQ** |
| 351 | **GWSATPIANG** | **SLGHDIHHPR** | GDAKK**YSQGN** | **VSAVGVTYDG** | **HTALTRVDWP** |
| 401 | **SAVVEGGSSG** | **SGLLTVAGDG** | **SYQLRGGLYG** | **GPSYCGAPTS** | **QRNDYFSDFS** |
| 451 | **GVYSQISR**YF | AP | | | |

**Figure S10: Sequence comparison of protein substrates of PIV studied here.**

Positions for which there is evidence of cleavage (and predicted cleavage at K81) are shown on the right, and sequence logo on the left compares residues neighbouring lysines in these sequences.

Cleaved sequences

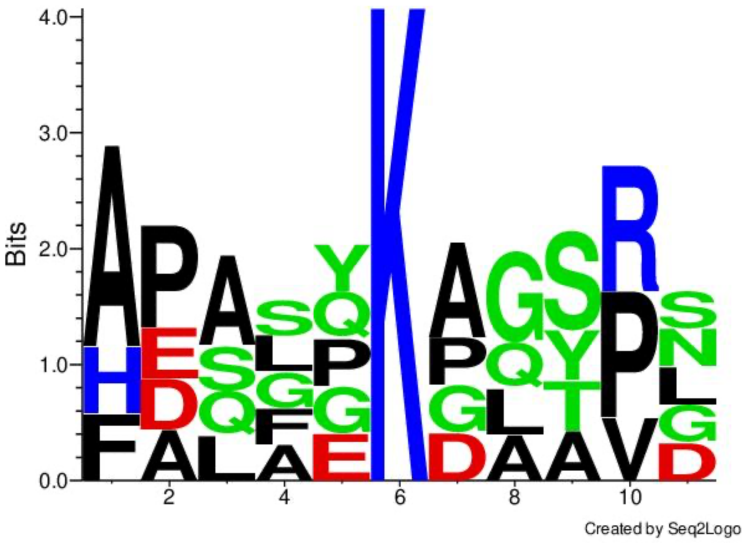

>Strep
HPQFEKGAAPG

>K81
APLAPKPGTPL
>K162
FEASGKDLSVN
>K211
ADSLYKAGYRD

>PaAP-K512
AAAGQKAQSRS

Uncleaved sequences

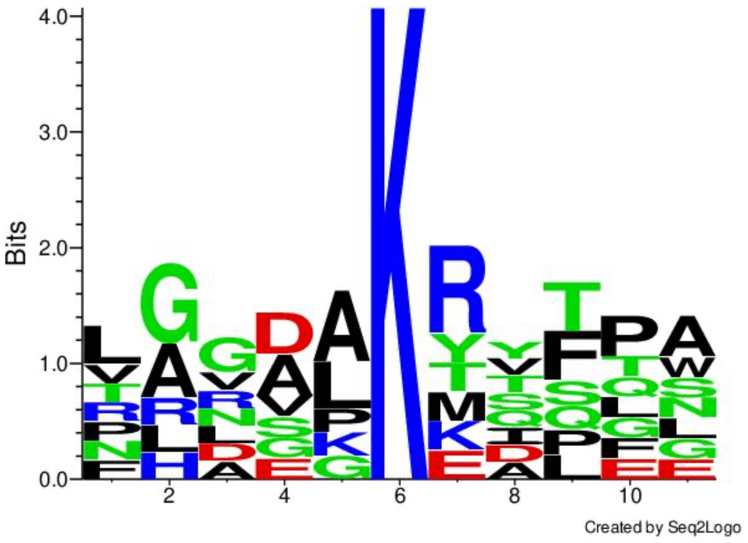

>K93
VGVGLKTATPE
>K154
FAGAGKEIFEA
>274
TAAVAKMVFTS
>K284
NGNSPKRQLFW
>K331
LHRDAKRDTLL
>K340
LLLELKRTPPA
>374
PRGDAKKYSQG
>K375
RGDAKKYSQGN

**Figure S11 – Mature PIV has 6 Lysine residues, majority are non-accessible for cleavage.**

Lysine residues are shown in pink, coloured by element. Active site residues are shown in yellow for reference.

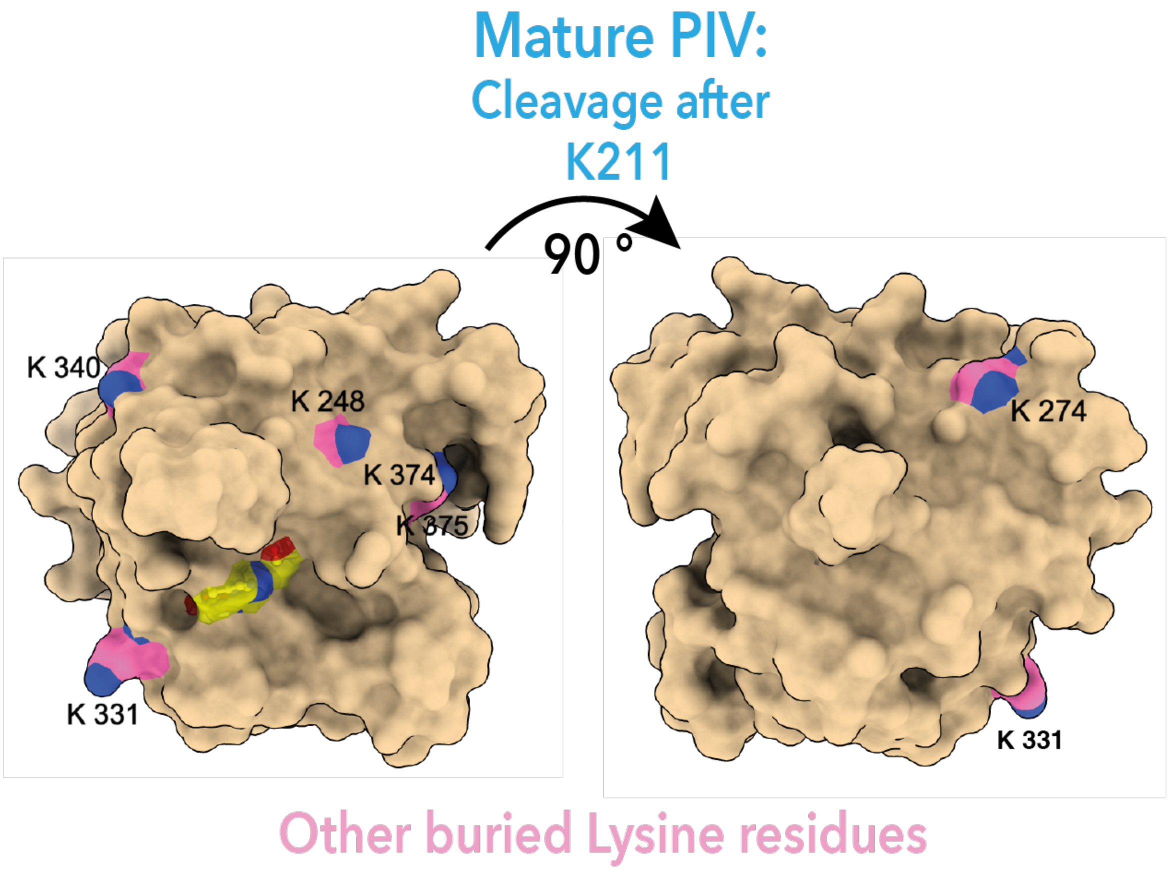

### Supplementary Information Tables

#### Supplementary Table 1. Primers

| | Primer | Sequence (5' to 3') | Purpose |
|---|---|---|---|
| **Recombinant Expression Constructs** | LysC_noSP_Insert_Fwd | TTTAAGGAGGTAAACATATGGCACCTGGCGCATCGGAG | Generate insert (secreted form of LysC lacking signal peptide) for Gibson assembly |
| | LysC_noSP_Insert_Rev | TGATGGTGGTGATGATGTTCAGGCGCGAAATAACGCG | |
| | LysC_noSP_Backbone_Fwd | TCTCGCGTTATTTCGCGCCTGAACATCATCACCACCATCATTG | Generate Backbone (pJ411) for Gibson assembly |
| | LysC_noSP_Backbone_Rev | ATCTCCGATGCGCCAGGTGCCATATGTTTACCTCCTTAAAAGTTAAAC | |
| **Mutagenesis** | LysC_S409A_Fwd | GGGTGGATCAgcTGGCTCGGGCCTTCTTAC | Generate S409A mutant in LysC noSP construct |
| | LysC_S409A_Rev | CCACTACGGCTGACGGC | |
| | LysC_strep_FWD | CAATTCGAGAAGGGAGCCGCACCTGGCGCATCG | *Generate Strep-tag at N-terminus of LysC noSP construct* |
| | LysC_strep_Rev | CGGATGTGACCACCCCATATGTTTACCTCCTTAAAAGTTAAACAAAATTATTTCTAG | |

**Supplementary Table 2. Plasmids and sequences**

| **Plasmids/ sequences** | **Description** |
|---|---|
| pJ411 | Recombinant gene expression with a non-cleavable C-terminal His8 tag. |
| Full length sequence of PIV | MHKRTYLNACLVLALAAGASQALAAPGASEMAGDVAVLQASPASTGHARFANPNAAISAAGI[illegible]<br>DGFGASGSCEVDAVCATQSGTRAYDNATAAVAKMVFTSSADGGSYICTGTLLNNGNSPKRQL[illegible]<br>GGLYGGPSYCGAPTSQRNDYFSDFSGVYSQISRYFAP |
| sequence of LysC_noSP_WT for protein expression | MAPGASEMAGDVAVLQASPASTGHARFANPNAAISAAGIHFAAPPARRVARAAPLAPKPGTPL[illegible]<br>AYDNATAAVAKMVFTSSADGGSYICTGTLLNNGNSPKRQLFWSAAHCIEDQATAATLQTIWFY[illegible]<br>DFSGVYSQISRYFAPEHHHHHHHH |
| sequence of LysC_Strep_WT for protein expression | MGWSHPQFEKGAAPGASEMAGDVAVLQASPASTGHARFANPNAAISAAGIHFAAPPARRVA[illegible]<br>VDAVCATQSGTRAYDNATAAVAKMVFTSSADGGSYICTGTLLNNGNSPKRQLFWSAAHCIED[illegible]<br>GAPTSQRNDYFSDFSGVYSQISRYFAPEHHHHHHHH |
| | KEY<br>Signal Peptide<br>His Tag<br>Strep Tag (WSHPQFEK)<br>Start Met |

#### Supplementary Table 3. Strains

| Strains | Description | Source |
|---|---|---|
| *E. coli* NEBa (DH5a) | *E. coli* cloning strain ( *fhuA2Δ(argF-lacZ)U169 phoA glnV44 Φ80Δ(lacZ)M15 gyrA96 recA1 relA1* ) | New England Biolabs (C2987) |
| *E. coli* SHuffle® T7 | *E. coli* expression strain ( F´ *lac, pro, lacI*$^q$ */ Δ(ara-leu)7697 araD139 fhuA2 lacZ::T7 gene1 Δ(phoA)PvuII phoR ahpC* galE (or U) galK λatt::pNEB3-r1-cDsbC* (Spec$^R$, *lacI*$^q$) *ΔtrxB rpsL150*(Str$^R$) *Δgor Δ(malF)3* ) | New England Biolabs (C3026J) |

#### Supplementary Table 4. Crystallisation conditions

| | Condition |
|---|---|
| **LysC S409A (Seed)** | **BCS G9**<br>0.1 M Magnesium Chloride hexahydrate, 0.1 M Sodium acetate trihydrate, 0.1 M Bis-Tris pH 6.5, 15 % v/v PEG Smear Broad<br><br>(PEG smear broad: 4.55%v/v PEG 400, 4.55%v/v PEG 500 MME, 4.55%v/v PEG 600, 4.55%w/v PEG 1000, 4.55%w/v PEG 2000, 4.55%w/v PEG 3350, 4.55%w/v PEG 4000, 4.55%w/v PEG 5000 MME, 4.55%w/v PEG 6000, 4.55%w/v PEG 8000, 4.55%w/v PEG 10000) |
| **LysC S409A** | **BCS D1**<br>0.2 M Ammonium Nitrate, 0.1 M Sodium cacodylate pH 5.3, 22.5 % v/v PEG Smear Low<br><br>(PEG Smear Low: 4.55%v/v PEG 400, 4.55%v/v PEG 500 MME, 4.55%v/v PEG 600, 4.55%w/v PEG 1000) |
| **Cryo-protectant** | Mother liquor + 20 % v/v Ethylene glycol |

**Supplementary Table 5. Crystallographic Data Table**

| | | LysC S409A |
|---|---|---|
| **Accession code** | | 31RM |
| **Data Collection** | | |
| | Resolution (Å) | 154.08 - 2.17 (2.21 - 2.17) |
| | Space group | C 2 2 21 |
| | Cell Dimensions a, b, c (Å) | 127.65, 209.33, 154.08 |
| | α, β, γ (°) | 90, 90, 90 |
| | Total reflections | 2531134 (55318) |
| | Unique reflections | 108486 (5152) |
| | Multiplicity | 23.3 (10.7) |
| | Completeness (%) | 99.7 (95.4) |
| | Mean I/sigma(I) | 12.6 (0.6) |
| | R-meas | 0.17 (2.34) |
| | R-pim | 0.034 (0.699) |
| | CC1/2 | 0.999 (0.315) |
| **Refinement** | | |
| | R-free | 0.201 |
| | R-work | 0.180 |
| | Total non-hydrogen atoms | 13026 |
| | Total macromolecule atoms | 12527 |
| | Total solvent atoms | 499 |
| | Protein molecules per ASU | 4 |
| | Residues per protein | 423 |
| | RMS(bonds) (Å) | 0.007 |
| | RMS(angles) (°) | 0.77 |
| | Ramachandran favoured (%) | 97.50 |
| | Ramachandran allowed (%) | 2.50 |
| | Ramachandran outliers (%) | 0 |
| | Average B-factor (Å2) | 60.1 |

*Values in parentheses are for the high-resolution shell
**R-value test set size =  5

**Supplementary Table 6. SIR targets for intact K- and O-peptides and their PIV generated cleavage fragments**

| | | Species | | | | |
|---|---|---|---|---|---|---|
| **Peptide** | **Fragment** | **$[M+H]^+$** | **$[M+2H]^{2+}$** | **$[M+3H]^{3+}$** | **$[M+Na]^+$** | **$[M+H+NA]^{2+}$** |
| **K-Peptide** | **Intact** APGTPLQVGVGLKTATPE | 1735.958829 | 868.483053 | 579.324460 | 1757.940771 | 879.474024 |
| | **N-Terminal Fragment** APGTPLQVGVGLK | 1236.731012 | 618.869144 | 412.915188 | 1258.712954 | 629.860115 |
| | **C-Terminal Fragment** TATPE | 518.245657 | 259.626467 | 173.420070 | 540.227599 | 270.617438 |
| **O-Peptide** | **Intact** APGTPLQVGVGL{ornithine}TATPE | 1721.943180 | 861.475228 | 574.652577 | 1743.925122 | 872.466199 |
| | **N-Terminal Fragment** APGTPLQVGVGL[ornithine} | 1222.715363 | 611.861320 | 408.243305 | 1244.697305 | 622.852291 |
| | **C-Terminal Fragment** TATPE | 518.245657 | 259.626467 | 173.420070 | 540.227599 | 270.617438 |
